# Maltotriose consumption by hybrid *Saccharomyces pastorianus* is heterotic and results from regulatory cross-talk between parental sub-genomes

**DOI:** 10.1101/679563

**Authors:** Nick Brouwers, Anja Brickwedde, Arthur R. Gorter de Vries, Marcel van den Broek, Susan M. Weening, Lieke van den Eijnden, Jasper A. Diderich, Feng-Yan Bai, Jack T. Pronk, Jean-Marc G. Daran

## Abstract

*S. pastorianus* strains are hybrids of *S. cerevisiae* and *S. eubayanus* that have been domesticated for several centuries in lager-beer brewing environments. As sequences and structures of *S. pastorianus* genomes are being resolved, molecular mechanisms and evolutionary origin of several industrially relevant phenotypes remain unknown. This study investigates how maltotriose metabolism, a key feature in brewing, may have arisen in early *S. eubayanus × S. cerevisiae* hybrids. To address this question, we generated a near-complete genome assembly of Himalayan *S. eubayanus* strains of the Holarctic subclade. This group of strains have been proposed to be the origin of the *S. eubayanus* subgenome of current *S. pastorianus* strains. The Himalayan *S. eubayanus* genomes harbored several copies of a *SeAGT1* α-oligoglucoside transporter gene with high sequence identity to genes encountered in *S. pastorianus*. Although Himalayan *S. eubayanus* strains are unable to grown on maltose and maltotriose, their maltose-hydrolase and *SeMALT1* and *SeAGT1* maltose-transporter genes complemented the corresponding null mutants of *S. cerevisiae*. Expression, in a Himalayan *S. eubayanus* strain, of a functional *S. cerevisiae* maltose-metabolism regulator gene (*MALx3*) enabled growth on oligoglucosides. The hypothesis that the maltotriose-positive phenotype in *S. pastorianus* is a result of heterosis was experimentally tested by constructing a *S. cerevisiae* × *S. eubayanus* laboratory hybrid with a complement of maltose-metabolism genes that resembles that of current *S. pastorianus* strains. The ability of this hybrid to consume maltotriose in brewer’s wort demonstrated regulatory cross talk between sub-genomes and thereby validated this hypothesis. These results provide experimental evidence of the evolutionary origin of an essential phenotype of lager-brewing strains and valuable knowledge for industrial exploitation of laboratory-made *S. pastorianus*-like hybrids.

**Importance:** *S.pastorianus*, a *S.cerevisiae* X *S.eubayanus* hybrid, is used for production of lager beer, the most produced alcoholic beverage worldwide It emerged by spontaneous hybridization and have colonized early lager-brewing processes. Despite accumulation and analysis of genome sequencing data of *S.pastorianus* parental genomes, the genetic blueprint of industrially relevant phenotypes remains unknown. Assimilation of wort abundant sugar maltotriose has been postulated to be inherited from *S.cerevisiae* parent. Here, we demonstrate that although Asian *S.eubayanus* isolates harbor a functional maltotriose transporter *SeAGT1* gene, they are unable to grow on α-oligoglucosides, but expression of *S. cerevisae* regulator *ScMAL13* was sufficient to restore growth on trisaccharides. We hypothesized that *S. pastorianus* maltotriose phenotype results from regulatory interaction between *S.cerevisae* maltose transcription activator and the promoter of *SeAGT1*. We experimentally confirmed the heterotic nature of the phenotype and thus this results provide experimental evidence of the evolutionary origin of an essential phenotype of lager-brewing strains.

## Introduction

*Saccharomyces pastorianus* is an interspecific hybrid of *S. cerevisiae* and *S. eubayanus* (1–4). *S. pastorianus* strains are widely used for production of lager beer, which is currently the most produced alcoholic beverage worldwide. Lager brewing requires alcoholic fermentation at relatively low temperatures. *S. pastorianus* was hypothesized to have emerged by spontaneous hybridization and to have colonized early lager-brewing processes due to a combination of cold-tolerance inherited from *S. eubayanus* and superior fermentation kinetics inherited from *S. cerevisiae* (5–7). Lager beer is brewed from barley wort, whose sugar composition consists, by weight, of approximately 15% glucose, 60% maltose and 25% maltotriose (8). During wort fermentation, maltotriose is generally only utilized after glucose and maltose are depleted, while its consumption is also relatively slow and often incomplete (9–11).

Complete sugar utilization is desirable for lager beer fermentation to optimize concentrations of ethanol and flavor compounds and to avoid residual sweetness (12). While *S. pastorianus* and *S. cerevisiae* strains are capable of consuming maltotriose, none of the wild isolates of *S. eubayanus* characterized thus far have been shown to possess this trait (6,13,14). These observations led to the hypothesis that the ability of *S. pastorianus* to ferment maltotriose was inherited from *S. cerevisiae* (6, 13, 15–17).

The genetic information for maltose utilization is well conserved in *Saccharomyces* species and depends on three gene families. *MALT* genes encode plasma-membrane proton symporters with varying substrate specificities and affinities (18, 19), *MALS* genes encode α-glucosidases that hydrolyze α-oligoglucosides into glucose, while *MALR* genes encode a regulator required for transcriptional induction of *MALT* and *MALS* genes by maltose (20, 21). In *Saccharomyces* species, maltose-utilization genes are generally organized in *MAL* loci. These loci contain a *MALT* gene (called *ScMALx1* and *SeMALTx* in *S. cerevisiae* and *S. eubayanus*, respectively), a *MALS* gene referred to as *ScMALx2* or *SeMALSx* and a *MALR* gene referred to as *ScMALx3* or *SeMALRx* (13, 22). In the absence of glucose and presence of maltose, the MalR regulator binds a bi-directional promoter, thereby simultaneously activating expression of *MALT* and *MALS* genes (23).

The *ScMAL1-ScMAL4* and *ScMAL6* loci of *S. cerevisiae* as well as the *SeMAL1-SeMAL4* loci of *S. eubayanus* are located in subtelomeric regions (13, 24–26). While all *S. cerevisiae* ScMalx1 transporters transport maltose, only ScMal11 is able to also transport maltotriose (9). *ScMAL11* (also known as *SeAGT1*) shares only 57% nucleotide identity with other *ScMALx1* genes (27). The four *SeMALT* (*SeMALT1-4*) genes identified in the genome of the Patagonian type strain FM1318/CBS 12357 of *S. eubayanus* were shown to encode functional maltose transporters, but none of these genes enabled maltotriose transport (13). While no clear *ScAGT1* ortholog was found in *S. eubayanus* CBS 12357^τ^, such an ortholog was recently found in the genomes of two North American isolates assigned to the Holarctic subclade of *S. eubayanus* (14).

*S. pastorianus* inherited *MAL* genes from both *S. cerevisiae* and *S. eubayanus* (2, 4, 28). However, the *S. cerevisiae-derived* maltotriose-transporter gene *ScAGT1* is truncated and, therefore, non-functional in *S. pastorianus* (10). Instead, maltotriose consumption by *S. pastorianus* strains was attributed to *SeAGT1* and *SpMTY1/SpMTT1* genes (29–32). In *S. pastorianus, SeAGT1* is located on *S. eubayanus* CHRXV and was therefore, already before the identification of an *AGT1* ortholog in Holarctic *S. eubayanus* strains (14), assumed to originate from *S. eubayanus* (2). *SpMTY1*, also referred to as *SpMTT1*, is located on *S. cerevisiae* CHRVII and has less than 92% sequence identity with other *Saccharomyces* maltose transporters (30). However, *SpMTY1* contains sequence patches with high similarity to maltose transporters from *S. eubayanus* and *S. paradoxus* (33). Recently, two independent laboratory evolution studies with *S. eubayanus* demonstrated that recombination of different *SeMALT* genes yielded chimeric, neo-functionalized genes that encoded maltotriose transporters (34, 35). *SpMTY1* may have resulted from successive introgressions of maltose-transporter genes from *S. cerevisiae, S. eubayanus* and *S. paradoxus*.

Recently made *S. cerevisiae × S. eubayanus* laboratory hybrids showed similar lager-brewing performance as *S. pastorianus* strains, also with respect to maltotriose utilization (6, 16, 17, 36). In these hybrids, maltotriose consumption depended on the presence of a functional *Sc*Agt1 transporter encoded by the *S. cerevisiae* subgenome (37). However, in view of the non-functionality of *ScAGT1* in current *S. pastorianus* strains, these laboratory hybrids did not fully recapitulate the genetic landscape of *S. pastorianus* with respect to maltotriose fermentation (2, 6, 36).

Studies on laboratory hybrids based on *S. eubayanus* strains whose genomes are more closely related to the *S. eubayanus* subgenome of *S. pastorianus* strains than that of the Patagonian type strain CBS12357 might generate new insights into the evolution of maltotriose utilization in *S. pastorianus*. To date, Himalayan *S. eubayanus* isolates show the highest sequence identity with the *S. eubayanus* sub-genome of *S. pastorianus*, with up to 99.82% identity, as opposed to 99.56% for *S. eubayanus* CBS12357^⊤^ (38).

Here, we investigated if and how the genomes of Himalayan *S. eubayanus* strains could have contributed to maltotriose utilization in the earliest hybrid ancestors of current *S. pastorianus* strains. To this end, we generated chromosome-level genome assemblies of these strains by long-read DNA sequencing. Since the Himalayan strains were unable to utilize maltotriose, we functionally characterized the assembled *MAL* genes and identified genetic determinants that prevented maltotriose utilization. Subsequently, a laboratory hybrid of a representative Himalayan *S. eubayanus* strain and a maltotriose-deficient ale strain of *S. cerevisiae* was generated to investigate the genetics of maltotriose utilization in a hybrid context. We discuss the implications of the experimental results for the proposed role and origin of *SeAGT1* in *S. pastorianus* and for the potential of hybridization to enable maltotriose consumption in novel *Saccharomyces* hybrids.

## Results

### Sequencing of Himalayan *S. eubayanus* strains revealed variations of sub-telomeric regions and presence of novel putative maltose transporter genes

It has been proposed that the *S. eubayanus* genetic pool of *S. pastorianus* was inherited from an ancestor of the Asian *S. eubayanus* lineage (38). With 99.82% identity, the Himalayan *S. eubayanus* strains CDFM21L.1 and ABFM5L.1 that belong to the Holarctic lineage (39), present the closest characterized relatives of the *S. eubayanus* ancestor of lager brewing yeasts. However, this distance was based on a limited sequencing space (38) and the analysis did not investigate the presence of specific *S. eubayanus* genetic markers found in *S. pastorianus* hybrids. Therefore, the genome of the Himalayan *S. eubayanus* strain CDFM21L.1 was sequenced with a combination of long-read and short-read techniques (Oxford Nanopore MinION and Illumina technologies, respectively) to generate a near-complete draft reference genome sequence. The resulting CDFM21L.1 genome assembly comprised 19 contigs, including its mitochondrial genome. All chromosomes were completely assembled from telomere to telomere, except for chromosome XII, which was fragmented into 3 contigs due to the repetitive rDNA region and manually assembled into a single scaffold. With a total size of 12,034,875 bp, this assembly represents the first near-complete draft genome of a *S. eubayanus* strain of the Holarctic clade (39).

Chromosome-level assemblies were hitherto only available for the Patagonia B-clade strain CBS 12357^⊤^ (1, 13). We identified three major structural differences in CDFM21L.1 relative to CBS12357^⊤^ using Mauve (40): (i) a paracentric inversion in the sub-telomeric region of chromosome VII, involving approximately 8 kbp, (ii) a translocation of approximately 12 kbp from the left sub-telomeric region of chromosome VIII to the right sub-telomeric region of chromosome VI, and (iii) a reciprocal translocation between approximately 20 kbp from the right sub-telomeric region of chromosome V and approximately 60 kbp from the center of chromosome XII (Figure 1A). All structural variation involved sub-telomeric regions, in accordance with their known relative instability (41–43).

**Figure 1.**
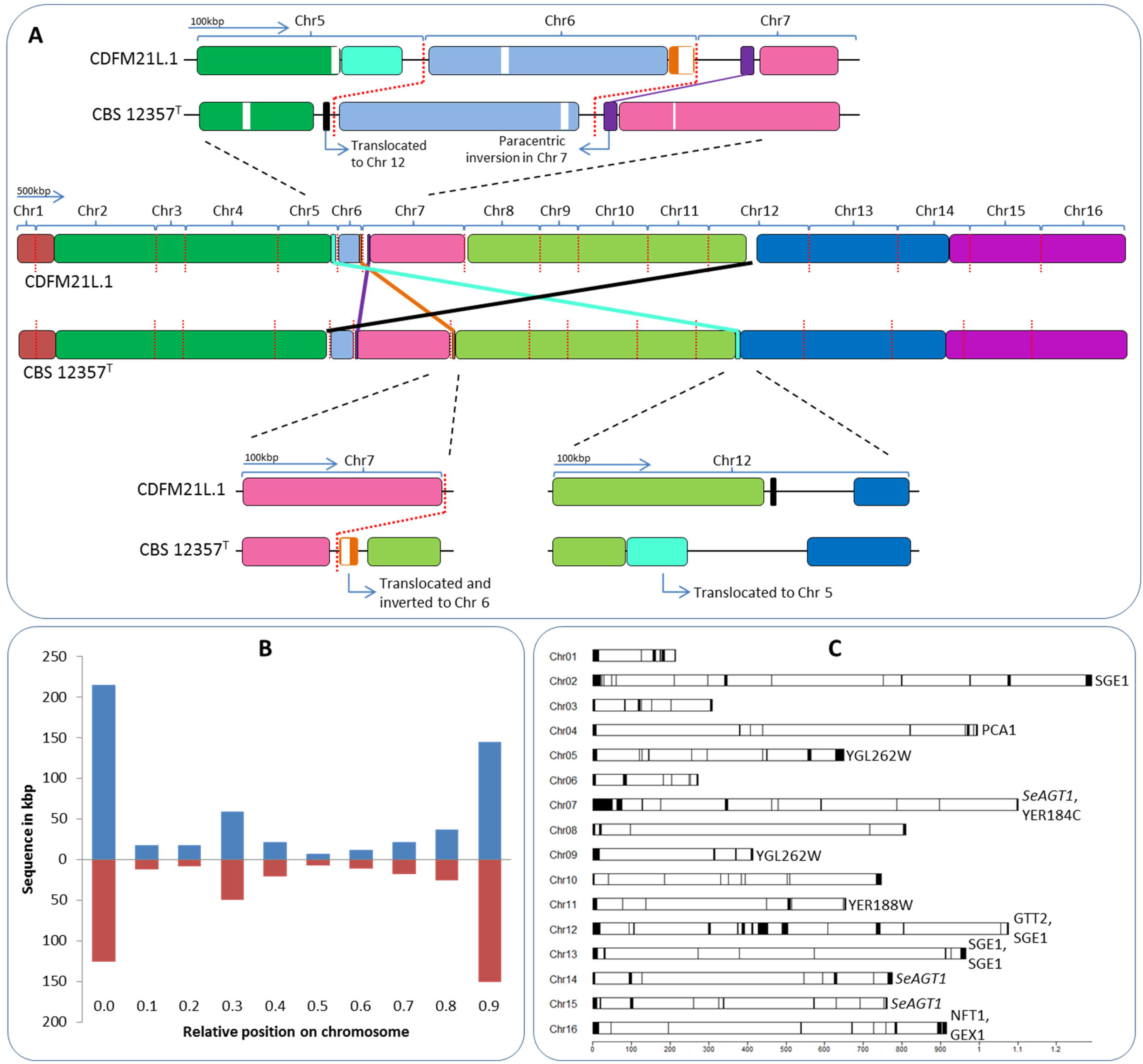
Genome comparison between CDFM21L.1 and CBS 12357^⊤^. **(A)** Translocations in CDFM21L.1 relative to CBS12357^⊤^, (i) a paracentric inversion in the sub-telomeric region of chromosome VII, involving approximately 8 kbp, (ii) a translocation of approximately 12 kbp from the left sub-telomeric region of chromosome VIII to the right sub-telomeric region of chromosome VI, and (iii) a reciprocal translocation between approximately 20 kbp from the right sub-telomeric region of chromosome V and approximately 60 kbp from the center of chromosome XII.Red lines depicted chromosomes separation. Genome syntheny is indicated with colored blocks. (B) Relative chromosome position of gene presence differences between CDFM21L.1 (blue) and CBS 12357^⊤^ (red). **(C)** Representation of the assembled CDFM21L.1 *S. eubayanus* chromosomes, the black boxes denote newly added sequences. New annotated open reading frames and gene entries modified relative to the CBS 12357T draft genome (13).

An alignment comparison of the CDFM21L.1 and CBS 12357^⊤^ genomes with MUMmer revealed that 557 kb were unique to CDFM21L.1 and reciprocally 428 kb were unique to CBS 12357^⊤^. Sequences unique to CBS 12357^⊤^ (3.6% of its genome) and to CDFM21L.1 (4.6% of its genome) were located primarily in sub-telomeric regions and in repetitive regions, such as rDNA on chromosome XII (Figure 1B). Out of the 32 sub-telomeric regions, 23 exhibited absence of synteny. Conserved synteny was observed for sub-telomeric regions on CHRIII (left), CHRIV (left and right), CHRVI (left), CHRIX (right), CHRXI (right), CHRXII (right), CHRXIV (left) and CHRXV (right) (Supplementary File 1).

The 428 kb of sequence that were absent in the Himalyan *S. eubayanus* strain included 99 annotated ORFs (Supplementary File 1). Of the 99 ORFs that were (partly) affected, 11 were completely absent in CDFM21L.1, involving mostly genes implicated in iron transport facilitation (Supplementary File 1). The 557 kb of sequence that was not present in CBS 12357^⊤^ included 113 annotated ORFs (Supplementary File 1). Of these 113 ORFs, 15 were completely absent in CBS 12357^⊤^. These 15 ORFs showed an overrepresentation of genes involved in transmembrane transport (Fishers exact test, P-value 4.8E^−5^) (Figure 1C).

Of the 15 ORFs unique to CDFM21L.1, three were identical orthologs of *S. cerevisiae MAL11/AGT1*. These three ORFs were found in the sub-telomeric regions of chromosomes VII, XIV and XV. Their sequence similarity with the *S. cerevisiae* CEN.PK113.7D and *S. pastorianus* CBS 1483 *MAL11/AGT1* genes was 82.7% and 99.89%, respectively. In addition to these *SeAGT1* genes, CDFM21L.1 genome sequence harbored three maltose transporters (*SeMALTx*), two maltases (*SeMALSx*) and three regulators (*SeMALRx*) encoding genes. In contrast to the situation in *S. eubayanus* CBS 12357^⊤^, none of the *SeMAL* genes formed a canonical *MAL* locus in CDFM21L.1 (Figure 2). A systematic sequence inspection of these CDFM21L.1 *SeMAL* genes revealed mutations that prematurely interrupted the reading frames of *SeMALR1* (CHRV), *SeMALT2* (CHRXII) and *SeMALT3* (CHRXII).

**Figure 2.**
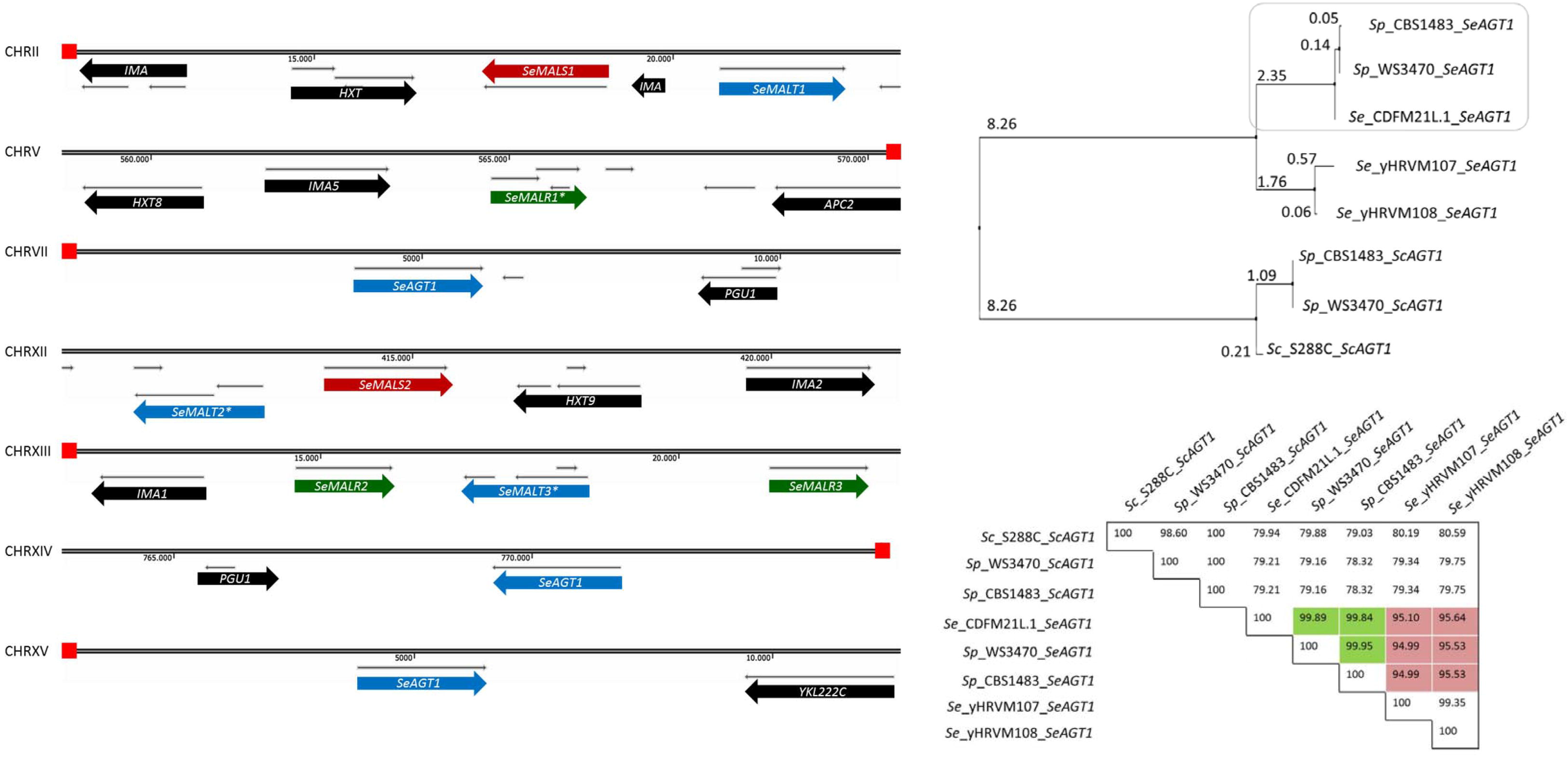
Organization of sub-telomeric regions involving *MAL* genes and *SeAGT1* in CDFM21L.1. **(A)** Chromosome sections are represented as lines and red boxes denote telomeres. CDFM21L.1 genome harbors three *SeMALT* genes of which *SeMALT2* and *SeMALT3* have a mutation resulting in an early stop codon and truncated protein (denoted with *). Three copies of *SeAGT1* were found close to the telomeres on CHRVII, XIV and XV. Furthermore there are two intact *SeMALS* genes on CHRII (1) and XII (2) and three *SeMALR* genes on CHRV (1) and XIII (2) whose copy on CHRV is also mutated (*SeMALR1*\*). The gene and interval sizes are approximately to scale. **(B)** Phylogeny of *Saccharomyces SeAGTl* genes described in *S. cerevisiae, S. eubayanus*, and lager-brewing hybrid *S. pastorianus*. (C) Nucleotide percentage identities between *AGT1* orthologs from *S. cerevisiae, S. eubayanus*, and lager-brewing hybrid *S. pastorianus*. Green color indicates highest similarity between SeAGT1 and SeAGT1 genes from *S. pastorianus* strains CBS 1483 and WS3470. Red color indicates similarity between SeAGTl from North American strains with SeAGT1 genes from Asian *S. eubayanus* and *S. pastorianus* strains CBS 1483 and WS3470.

In addition to *S. eubayanus* CDFM21L.1 strain, a second Himalayan *S. eubayanus* isolate (ABFM5L.1) was sequenced. These two strains were 99.97% genetically identical at the nucleotide level, their *MAL* genes were syntenic and the premature stop codons in *SeMALR1* (CHRV), *SeMALT2* (CHRXII) and *SeMALT3* (CHRXII) were conserved. Two additional mutations were identified in one of the three *SeAGT1* genes. A nucleotide variation on position 53 and 939 (T instead of an A and A instead of a G)) resulted in a glycine to valine and arginine to lysine change, respectively.

### Paradoxically, Himalayan *S. eubayanus* strains do not utilize maltose and maltotriose

Identification of *SeAGT1* in the two Himalayan *S. eubayanus* strains suggests an ability to not only grow on maltose but also on maltotriose. Strains from the Holarctic clade have previously been hypothesized to be the donor of the *S. eubayanus* sub-genome in *S. pastorianus* hybrids (38, 39). However, no physiological data regarding their ability to grow on the sugars present in wort are available. To assess their growth characteristics, the *Asian S. eubayanus* strains CDFM21L.1 and ABFM5L.1, the Patagonian *S. eubayanus* type strain CBS 12357^⊤^ and the *S. pastorianus* strain CBS 1483 were grown on diluted industrial brewer’s wort at 12 °C. As reported previously, *S. pastorianus* strain CBS 1483 could utilize all three sugars but did not fully consume maltotriose (Figure 3) (11). Also in accordance with previous observations (6), CBS 12357^⊤^ consumed glucose and maltose completely, but left maltotriose untouched. However, in marked contrast to *S. eubayanus* CBS 12357^⊤^, neither CDFM21L.1 nor ABFM5L.1 consume maltose after growth on glucose. Moreover, like CBS 12357^⊤^, maltotriose was not metabolized by these two *S. eubayanus* strains. While in CBS 12357^⊤^ an ability to grow on maltose and an inability to grow on maltotriose could be readily attributed to its *MAL* genes complement, CDFM21L.1 and ABFM5L.1 failed to grow on maltose even though they appeared to contain complete genes encoding maltose (*SeMALT1* and *SeAGT1*) and maltotriose (*SeAGT1*) transporters.

**Figure 3.**
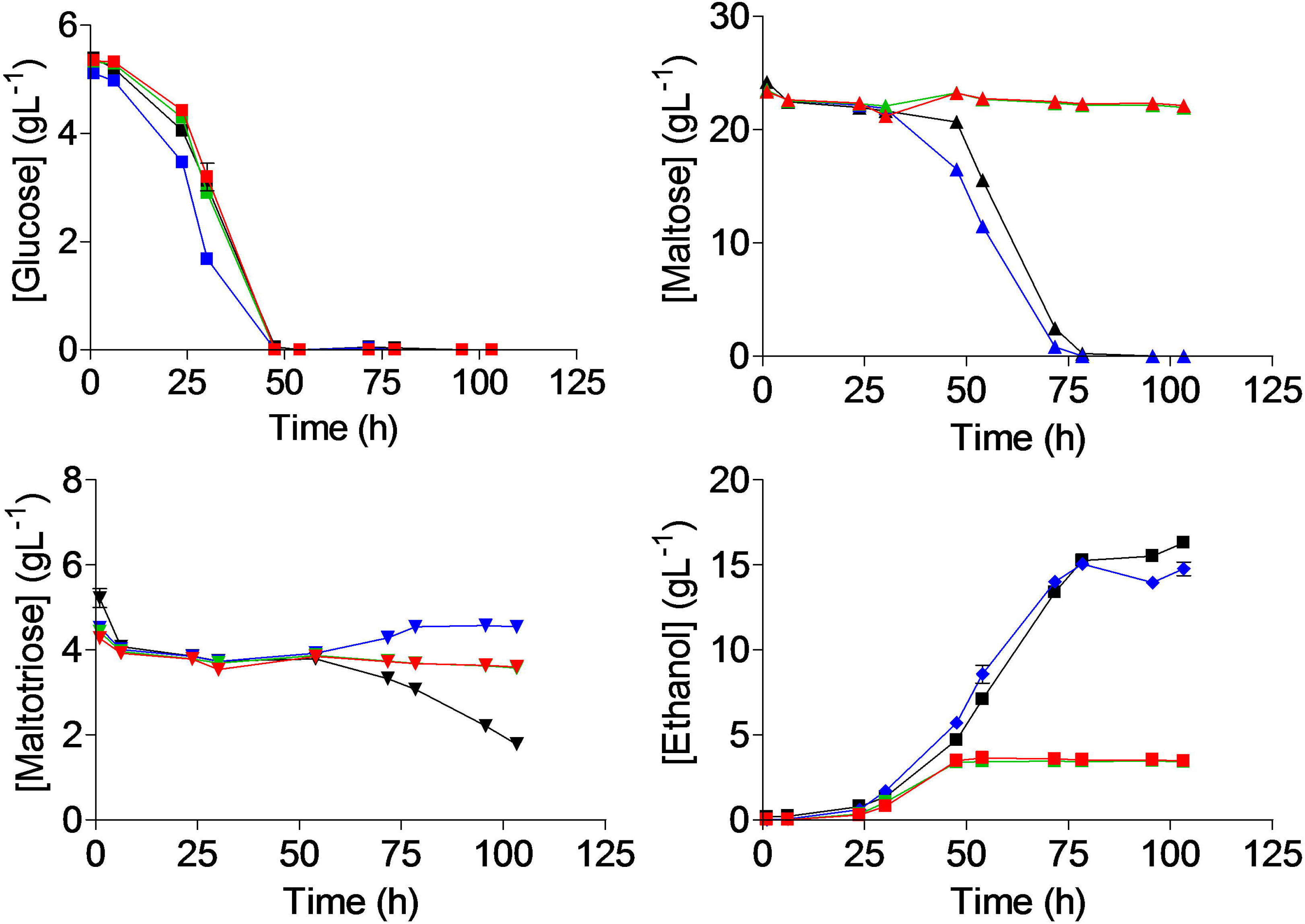
Characterization of sugar consumption of *S. pastorianus* CBS1483 (black) and *S. eubayanus* CBS 12357^⊤^ (blue), CDFM21L.1 (red), and ABFM5L.1 (green) on wort. For every sample, glucose (■), maltose (▲), maltotriose (▼) and ethanol (♦) were measured from the supernatant. Strains were grown at 12 °C for 110 hours in infusion Neubor flasks. Samples were filtered through a 0.22 μm filter and analyzed on HPLC. Data represents average and standard deviation of three biological replicates.

### Growth defects on maltose and maltotriose are caused by deficiency of the regulatory *Se*MalR proteins in *S. eubayanus* CDFM21L.1

The recent characterization of maltose metabolism in CBS 12357^⊤^ showed that the coding regions of transcriptionally silent maltose-transporter genes in *S. eubayanus* can potentially encode functional proteins (13). The inability of the *S. eubayanus* Himalayan isolates to grow on α-oligosaccharides precluded direct testing of transporter-gene functionality by deletion studies. Instead, these genes were expressed in *S. cerevisiae* IMZ616, which is devoid of all native maltose metabolism genes (44). The CDFM21L.1 transporter genes *SeMALT1, SeMALT2, SeMALT3* or *SeAGT1* were integrated at the *ScSGA1* locus in IMZ616 along with the *S. cerevisiae* maltase gene *ScMAL12* (13), yielding a series of strains overexpressing a single transporter (IMX1702 (*SeMALT1*), IMX1704 (*SeMALT2*), IMX1706 (*SeMALT3*) and IMX1708 (*SeAGT1*)). These strains, as well as the negative and positive control strains IMZ616 and IMX1365 (IMZ616 expressing *ScAGT1* and *ScMAL12*), were grown on SM media supplemented with either maltose or maltotriose. On maltose, not only the positive control strain IMX1365, but also IMX1702 (*SeMALT1*) and IMX1708 (*SeAGT1*) were able to grow on maltose, consuming 30 and 60%, respectively, of the initially present maltose after 100 h (Figure 4A). As anticipated, the *SeMALT2* and *SeMALT3* alleles with premature stop codons did not support growth on maltose. Of the two strains that grew on maltose, only IMX1708 (*SeAGT1*) also grew on maltotriose. These results demonstrate that *SeAGT1* from a Holarctic *S. eubayanus* encoded a functional maltotriose transporter and, consequently, that the inability of Holarctic strains to grow on maltose and maltotriose was not caused by transporter dysfunctionality.

**Figure 4.**
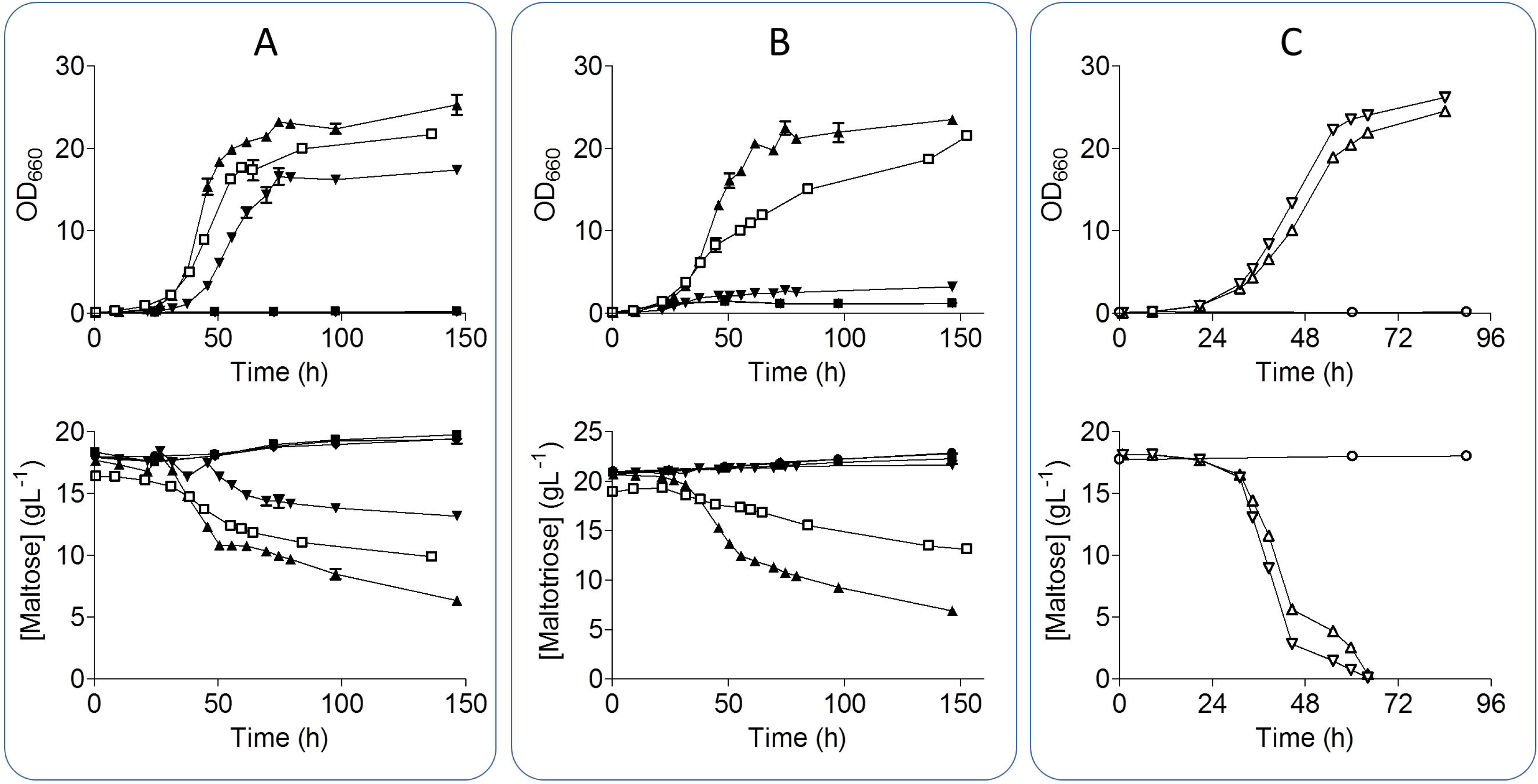
Overexpression of *SeMALT, SeAGT1* and *SeMALS* genes in a maltose negative background *S. cerevisiae* strain. Maltose negative background strain IMZ616 (■), IMX1365 overexpressing *ScMAL11* (▲), IMX1702 overexpressing *SeMALT1* (▼), IMX1704 overexpressing *SeMALT2* (♦) IMX1706 overexpressing *SeMALT3* (◎) and IMX1708 overexpressing SeAGT1 (□) were grown on SM 2% maltose or maltotriose at 20 °C. Growth on maltose **(A)** and on maltotriose **(B)** was monitored based on optical density (OD_660nm_) and concentrations of maltose and maltotriose in culture supernatants were measured by HPLC. Data are presented as average and standard deviation of two biological replicates. **(C)** IMX1313 overexpressing only *ScMAL31* (●), IMZ752 overexpressing *ScMAL31* and *SeMALS1* (△) and IMZ753 overexpressing *ScMAL31* and *SeMALS2* (▽) grown on SM maltose 2%. Growth was monitored based on optical density measurement at 660 nm (OD_660nm_) and maltose in culture supernatants was measured by HPLC. Data represents average and standard deviation of two biological replicates.

In addition to transport, metabolism of α-oligoglucosides requires maltase activity. Functionality of the putative *SeMALS1* and *SeMALS2* maltase genes was tested by constitutive expression in strain IMZ616, together with a functional ScMAL31 transporter genes, yielding strains IMZ752 and IMZ753, respectively. The maltase-negative strain IMX1313 was used as negative control. In SM medium with maltose, both IMZ752 (*SeMALS1*) and IMZ753 (*SeMALS2*) grew and completely consumed maltose within 65 h, demonstrating functionality of both hydrolase genes (Figure 4C).

In *S. cerevisiae* transcriptional regulation of *MALx2* and *MALx1* genes is tightly controlled by a transcription factor encoded by *MALx3* genes. Malx3 binds an activating site located in the bidirectional promoters that control expression of *MALx2* and *MALx1* genes (45, 46). To test whether absence of maltose consumption in Himalayan *S. eubayanus* strains was caused by a lack transcriptional upregulation of *SeMALT* and *SeMALS*, the *S. cerevisiae ScMAL13* gene was integrated at the *SeSGA1* locus in *S. eubayanus* CDFM21L.1, under the control of a constitutive *ScPGK1* promoter and *ScTEF2* terminator. *ScMAL13* expression in CDFM21L.1 enabled growth on maltose and maltotriose (Figure 5A), indicating that a lack of transcriptional upregulation was indeed the cause of the parental strain’s inability to grow on these oligoglucosides. However, consumption of maltose and maltotriose was incomplete and consumed sugars were almost exclusively respired, as no ethanol was measured after 60 h of cultivation.

**Figure 5.**
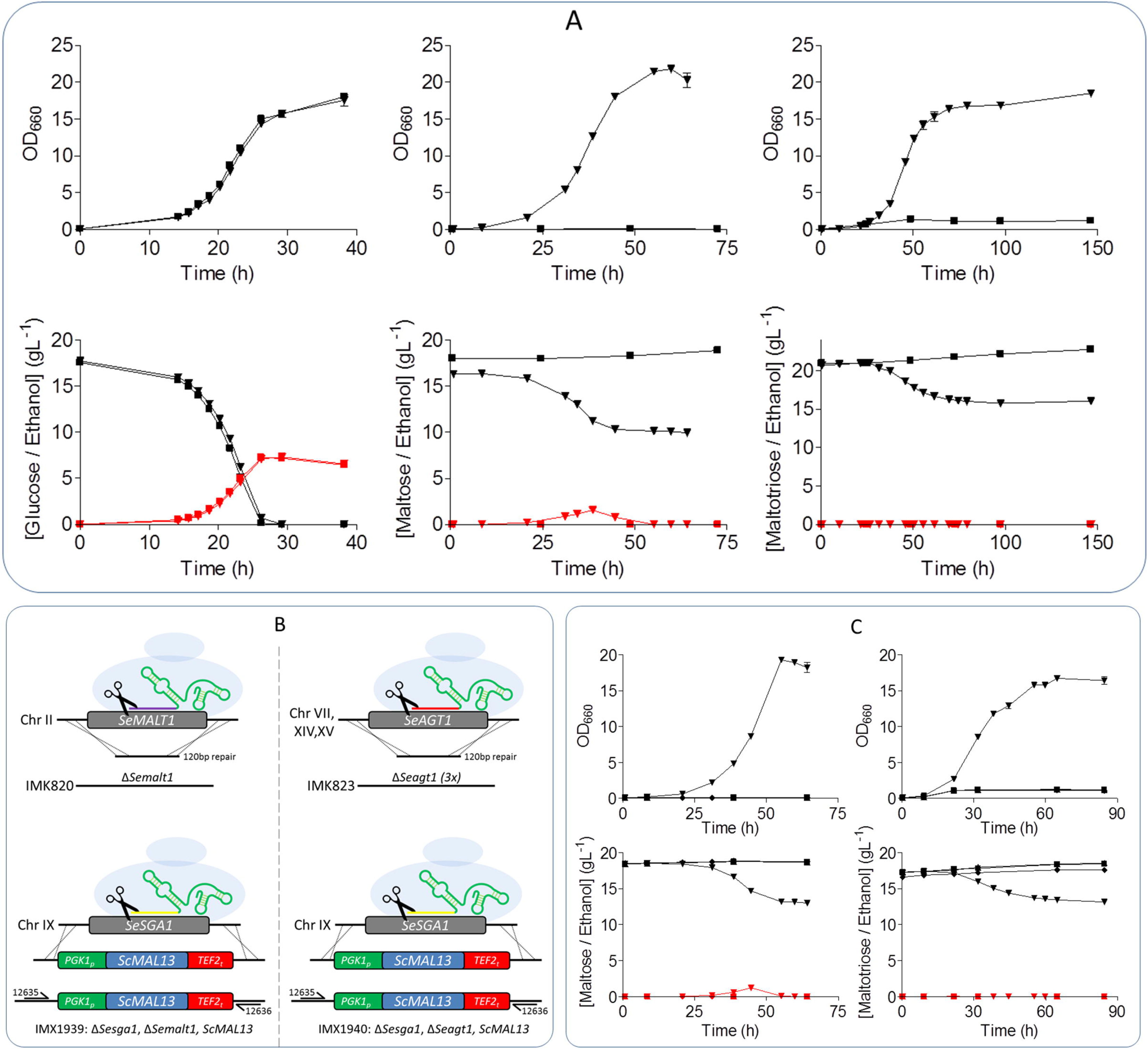
Integration of *ScMAL13* in CDFM21L.1 restores growth on maltose and maltotriose and enables native *SeMALT1* and *SeAGT1* characterization in knockout strains IMK820 and IMK823. **(A)** Characterization of *S. eubayanus* IMX1765 overexpressing *ScMAL13* (▼) and CDFM21L.1 (■) on SM with glucose, maltose or maltotriose at 20 °C. OD_660nm_ was measured (black) and sugar (black) and ethanol (red) concentrations were determined from the supernatant by HPLC. **(B)** Overview of constructed knockout strains. Knockouts of *SeMALT1* (IMK820) and *SeAGT1* (IMK823) were made with CRISPR-Cas9. Subsequently the *SeSGA1* locus was replaced by *ScPGK1_p_-ScMAL13-ScTEF2_t_* using CRISPR-Cas9 in both strains resulting in IMX1939 and IMX1940, respectively. **(C)** *S. eubayanus* strains IMK820 (■), IMK823 (▲), IMX1939 (▼) and IMX1940 (♦) were characterized on SM with maltose or maltotriose at 20 °C. OD_660nm_ was measured (black) and sugar (black) and ethanol (red) concentrations were determined from the supernatant by HPLC. All data represents average and standard deviation of biological duplicates.

The possibility to grow an engineered variant of *S. eubayanus* CDFM21L.1 on α-oligoglucosides offered an opportunity to study transporter function in its native context. Complementary functional characterization by gene deletion of *SeMALT1* and *SeAGT1* was performed using CRISPR-Cas9 genome editing method (13, 47). Deletion of *SeMALT1* and *SeAGT1* in CDFM21L.1 resulted in strains IMK820 and IMK823, respectively. Complete deletion of *SeAGT1* required disruption of six alleles. To confirm the complete removal of all copies, the genome of IMK823 was sequenced. Mapping reads onto the reference *S. eubayanus* CDFM21L.1 genome assembly confirmed that all six alleles were removed simultaneously. Subsequently, the regulator expression cassette (*ScPGK1_p_-ScMAL13-ScTEF2_t_*) was integrated in IMK820 and IMK823 at the *SeSGA1* locus yielding strains IMX1939 and IMX1940, respectively (Figure 5B). The four deletion strains IMK820(*SemalT1*Δ), IMK823 (*Seagt1*Δ), IMX1939 (*SemalT1*Δ *Sesga1*Δ::*ScMAL13*) and IMX1940 (*Seagt1*Δ *Sesga1*Δ::*ScMAL13*) were characterized on SMG, SMM or SMMt. All four strains were able to grow on glucose (Supplementary Figure 3). While strains IMK820, IMK823 and IMX1940 were unable to grow on maltose or maltotriose (Figure 5C), strain IMX1939 (*SemalT1*Δ *Sesga1*Δ::*SeMAL13*), which harbored functional *SeAGT1* copies, grew on maltose as well as on maltotriose. However, after 64 h of growth, these sugars were only partially consumed. Only 1.2 g L^−1^ ethanol was produced from maltose and no ethanol formation was observed during growth on maltotriose. The low ethanol concentration and the relatively high OD_660nm_ suggest that, under the experimental conditions strain IMX1939 exhibited a Crabtree negative phenotype and exclusively respired maltotriose. *S. eubayanus* IMX1940 (*Seagt1*Δ *Sesga1*Δ::*SeMAL13*) did not consume maltotriose after 84 h of incubation. Moreover, despite the presence of *SeMALT1*, which encoded a functional maltose transporter upon expression in *S. cerevisiae* IMZ616, strain IMX1940 was also unable to consume maltose.

In addition to a functional Malx3 transcription factor, transcriptional activation of *MAL* genes also requires presence of a *cis*-regulatory motif in the promoter of regulated genes. Transcriptome analysis of *S. eubayanus* CBS 12357^⊤^ recently showed that absence of a canonical *cis*-regulatory motif in *SeMALT1* and *SeMALT3* of *S. eubayanus* CBS 12357^⊤^caused a deficiency in their expression (13). To further explore regulation of *SeMAL* and *SeAGT1* genes, we investigated the impact of carbon sources on genome-wide transcriptome and, specifically, on transcriptional activation of genes involved in maltose metabolism. Duplicate cultures of *S. eubayanus* strain IMX1765 (*ScPGK1*_p_-*ScMAL13-ScTEF2*_t_) were grown on SMG, SMM and SMMt at 20 °C and sampled in mid-exponential phase. After mRNA isolation and processing, cDNA libraries and reads were assembled onto the newly annotated *S. eubayanus* CDFM21L.1 genome to calculate FPKM (fragments per kilobase of feature (gene) per million reads mapped) expression values. The heterologous regulator *ScMAL13*, expressed from the constitutive *ScPGK1* promoter, displayed the same expression in glucose- and maltose-grown cultures. Although *ScMAL13* was efficiently expressed on glucose, none of the nine *S. eubayanus* maltose genes (the three identical *SeAGT1* copies being undistinguishable) were transcriptionally induced under these conditions (Figure 6), confirming that the hierarchical regulatory role of glucose catabolite repression (45, 48) also takes place in *S. eubayanus*. During growth on maltose, all nine genes were significantly upregulated relative to glucose-grown cultures but large variations in expression level were observed. The maltase genes *SeMALS1* and *SeMALS2* and the transporter gene *SeAGT1* showed the highest upregulation, with fold changes of 148, 161 and 2355 respectively. Although upregulated *SeMALT1* displayed a fold change of 13, its normalized expression in maltose-grown cultures was 886-fold lower than that of *SeAGT1*. This weaker upregulation might explain why, despite the ability of its coding region to support synthesis of a functional maltose transporter, *SeMALT1* alone could not restore growth on maltose. The transcriptome data also revealed that the absence of maltose induction in CDFM21L.1 was not associated with defective *cis*-regulatory elements in *SeMALR* promoter sequences since the regulator genes were properly activated, instead these results would suggest that the SeMalR regulator are not functional.

**Figure 6.**
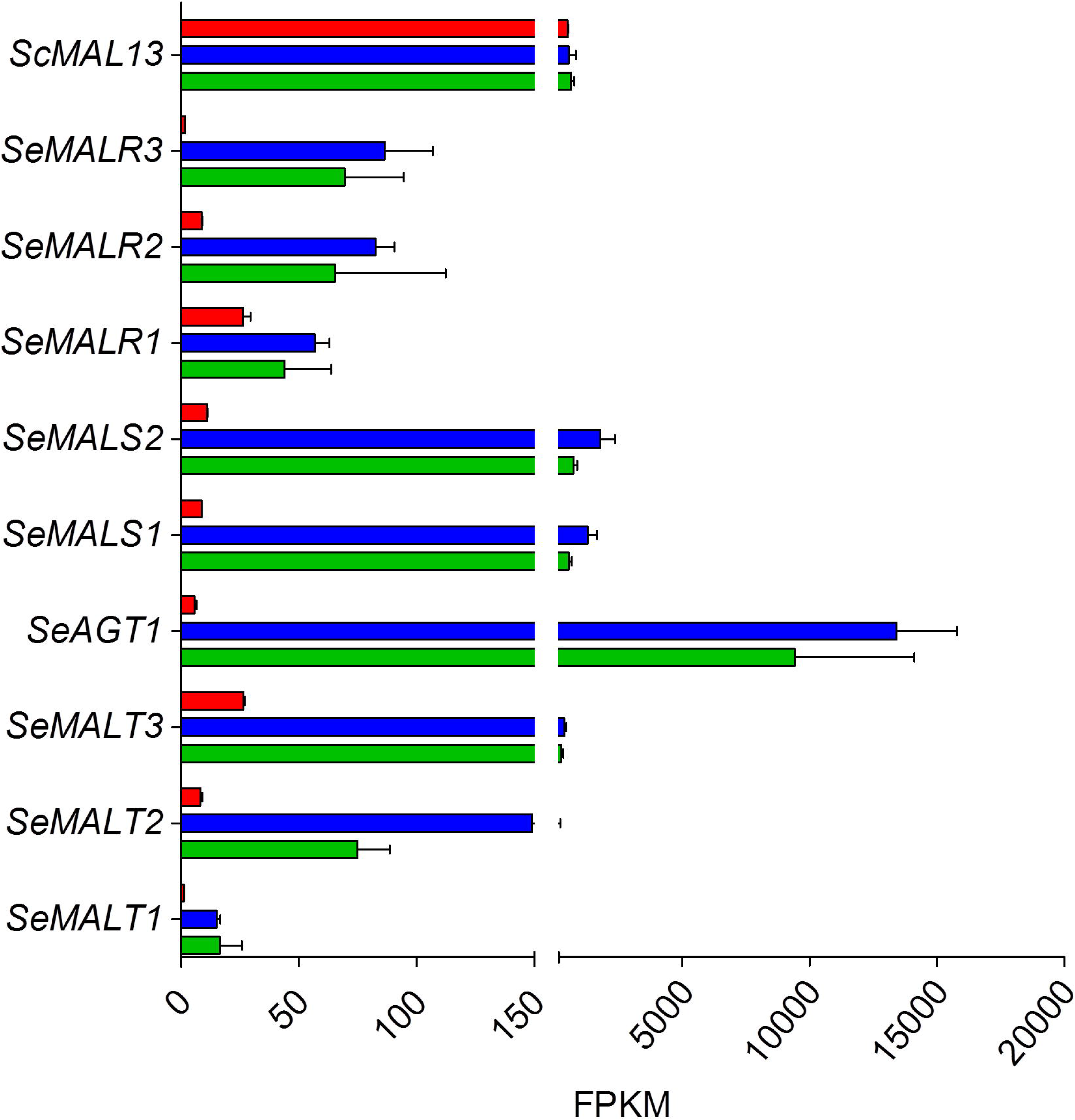
Expression levels of maltose metabolism genes in IMX1765. Normalized transcript levels of maltose metabolism genes from IMX1765mid-exponential phase grown on glucose (red), maltose (blue)and maltotriose (green) at 20 °C were calculated from duplicate RNA sequencing experiments (2x 150 bp) using the FPKM method. All data represents average and standard deviation of two biological duplicates.

### Hybridization of two maltotriose-deficient *S. eubayanus* and *S. cerevisiae* lineages results in heterosis through regulatory crosstalk

The genetic make-up of *S. pastorianus* lager-brewing yeasts strongly advocates that they originate from hybridization of *S. cerevisiae* and *S. eubayanus* parental lineages that were both unable to metabolize maltotriose (2). This hypothesis is consistent with the recurrent mutation in the *S. cerevisiae AGT1* allele of *S. pastorianus* strains as well as with the inability of Himalayan strains of *S. eubayanus* to grow on these oligoglucosides.

Spores of the Himalayan *S. eubayanus* CDFM21L.1 were hybridized with *S. cerevisiae* CBC-1. This top-fermenting *S. cerevisiae* is recommended for cask and bottle conditioning and unable to consume maltotriose (Lallemand, Montreal, Canada). Analysis of the CBC-1 assembly, obtained by a combination of long and short read sequencing, linked its maltotriose-negative phenotype to a total absence of the *MAL11/AGT1* gene. The resulting laboratory interspecific hybrid HTSH020 was characterized at 12 °C on synthetic wort, a defined medium whose composition resembles that of brewer’s wort. While *S. eubayanus* CDFM21L.1 only consumed glucose and *S. cerevisiae* CBC-1 consumed glucose and maltose after 103 h (Figure 7A), the interspecific hybrid HTSH020 completely consumed glucose, maltose and partially consumed maltotriose after 105 h, thus resembling characteristics of *S. pastorianus* strains (e.g. CBS1483 (11)). In addition to this gain of function, the hybrid HTSH020 outperformed both its parents on maltose consumption, since it depleted this sugar in 70 h instead of 95 h for strain CBC-1. Since *S. cerevisiae* grows generally slower at 12 °C, the experiments were also performed at 20 °C where HTSH020 consumed all maltose 16 h earlier than CBC-1 (Supplementary Figure 4).

**Figure 7.**
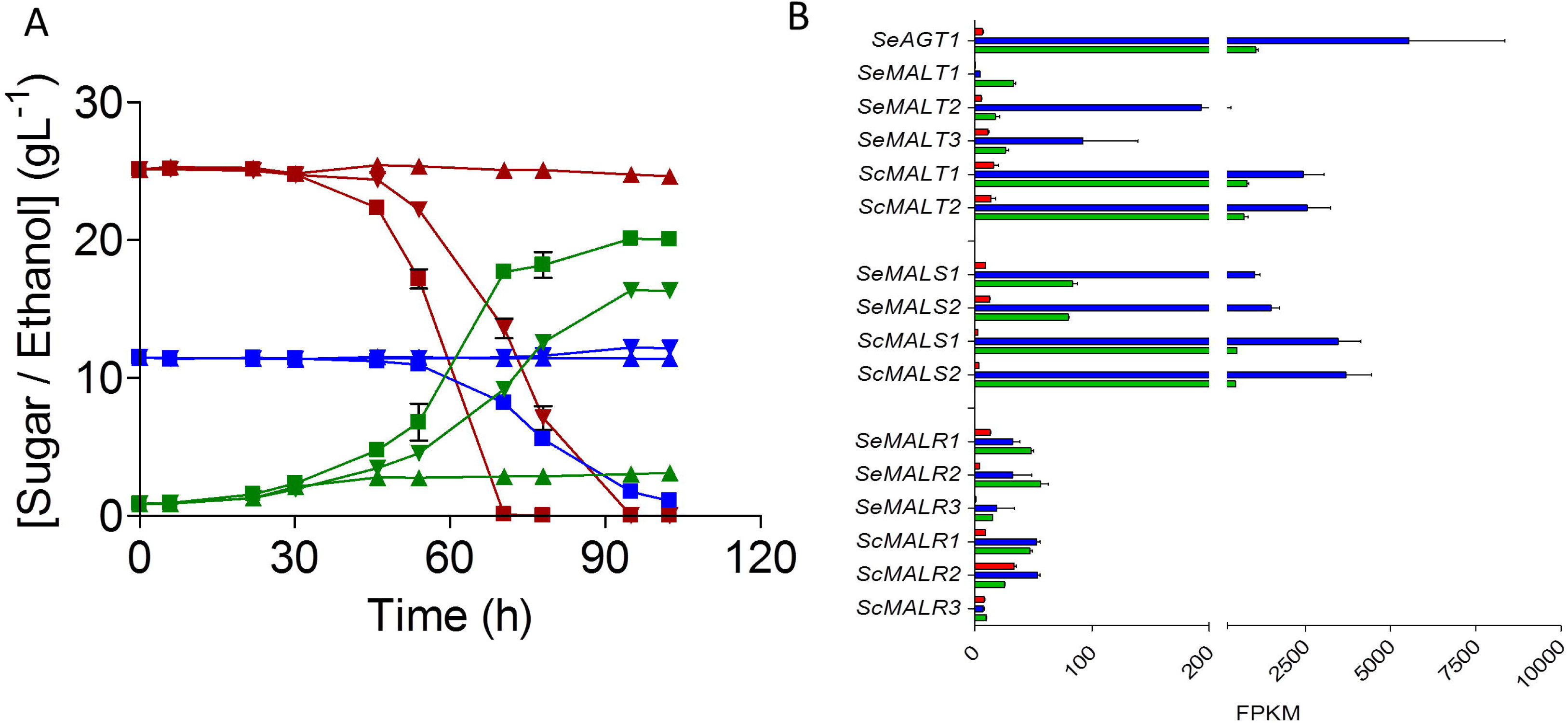
Hybridization of maltotriose deficient *S. cerevisiae* and *S. eubayanus* leading to crosstalk restoring maltotriose utilization, explains *S. pastorianus* phenotype. **(A)** Characterization of *S. cerevisiae* CBC-1 (▼), *S*. eubayanus CDFM21L.1 (▲) and hybrid HTS020 (■) on mock wort at 12 °C. Consumption of maltose (red), maltotriose (blue) and production of ethanol (green) was measured from supernatant by HPLC. Data represents average and standard deviation from biological triplicates. **(B)** Normalized transcript levels of maltose metabolism genes from HTS020 mid-exponential phase grown on glucose (red), maltose (blue)and maltotriose (green) at 20 °C were calculated from duplicate RNA sequencing experiments (2x 150 bp) using the FPKM method. All data represents average and standard deviation of two biological duplicates.

Transcriptome analysis of the hybrid strain HTSH020 grown on SM with different carbon sources showed that *SeAGT1* expression was repressed during growth on glucose, with a normalized expression level of 7 FPKM (Figure 7B). When grown on SM maltose, *SeAGT1, SeMALS1* and *SeMALS2* were significantly induced, with fold increases of 816,109 and 116, respectively (Figure 7B). Although *SeMALT1* and *SeMALT2* were induced, these transporters do not contribute to maltose metabolism due to truncation of their ORFs. These transcriptome data demonstrated that *SeAGT1* and *SeMALS* genes are induced by regulatory crosstalk between regulators encoded from the CBC-1 *S. cerevisiae* sub-genome and maltotriose transporter genes harbored by the *S. eubayanus* genome. This laboratory hybridization experiment may be the closest reproduction yet of how, centuries ago, maltotriose-fermentation capacity arose in the first hybrid ancestor of *S. pastorianus*.

## Discussion

The ability to consume maltose and maltotriose represents a key performance indicator of *S. pastorianus* lager-brewing strains (10). This study demonstrates how mating of *S. cerevisiae* and *S. eubayanus* strains that cannot themselves ferment maltotriose, can yield maltotriose-fermenting hybrids. This laboratory study illustrates how, centuries ago, maltotriose-fermentation capacity may have arisen in the first hybrid ancestor of *S. pastorianus*.

While the origin of the *S. eubayanus* parent of *S. pastorianus* strains is still under debate (49–51), phylogenetic analysis suggested a Far East Asian origin (38). However, this interpretation was based on a limited sequencing space and was constrained by the quality of available sequence assemblies. Since an ortholog of *SeAGT1* had previously only been found in the *S. eubayanus* sub-genome of *S. pastorianus* strains, this finding revived the discussion on the geographical origin of the ancestral *S. eubayanus* parent (14). The high-quality, annotated genome assemblies of the Himalayan *S. eubayanus* strains CDFM21L.1 and ABFM5L.1 presented in the present study revealed several copies of *SeAGT1*, whose very high sequence identity with *S. pastorianus SeAGT1* are consistent with the previously proposed Asian origin of the *S. eubayanus* sub-genome of *S. pastorianus* (14, 38, 39). Next, genome-sequence comparison of the Patagonian B sub-clade *S. eubayanus* strain CBS12357^⊤^ and the Holarctic sub-clade strain CDFM21L.1 revealed homoplasy of *SeAGT1*, probably reflecting that these sub-clades evolved in different ecological niches.

The *S. eubayanus* wild stock whose genome sequence most closely corresponds to the *S. eubayanus* sub-genome of *S. pastorianus* originates from the Tibetan plateau of the Himalaya (38). However, the first *S. cerevisiae x S. eubayanus* hybrid, from which current lager yeasts evolved by centuries of domestication, likely originates from a region between Bavaria and Bohemia in Central Europe. So far, European *S. eubayanus* isolates have not been reported. This may indicate that the original hybridization event occurred elsewhere or that the ancestral European lineage went extinct. The recent detection, in a metagenomics analysis of samples from the Italian Alps, of ITS1 sequences corresponding to *S. eubayanus* could indicate that a wild European lineage exists after all (52).

Functional characterization by heterologous complementation of an *S. cerevisiae* mutant strain established that the *Se*Agt1 transporters from the Himalayan *S. eubayanus* strains CDFM21L.1 and ABFM5L supported uptake of maltose and maltotriose. After showing that these strains also encoded a functional maltase gene, their inability to grow on maltose and maltotriose was attributed to an inability to transcriptionally upregulate maltose metabolism genes likely caused by loss of function mutations the regulator. In *S. cerevisiae* and, to some extent, in *S. eubayanus* strains of the Patagonian B sub-clade such as CBS12357^⊤^ (13, 26, 53), MAL loci exhibit a specific organization in which a transporter (*MALT*) and a hydrolase (*MALS*) gene are expressed from the same bidirectional promoter and are located adjacent to a regulator gene (*MALR*) (21). In contrast, of the seven genomic regions harboring *MAL* genes in the two Asian *S. eubayanus* strains, none showed this canonical organization (Figure 2) and the subtelomeric regions carrying *SeAGT1* did not harbour sequences similar to hydrolase or regulator genes. Subtelomeric regions harboring the other *MAL* genes indicated intensive reorganization as a result of recombination. In particular, subtelomeric regions on CHRII, CHRV and CHRXII provide clear indications for recombination events that scattered genes from ancestral MAL1 and MAL2 loci over several chromosomes. A similar interpretation could explain the reorganization MAL3 on CHRXIII (Figure 2). Similar events may have contributed to loss of function of the MAL regulators (MalR), as exemplified by the occurrence of a non-synonymous mutation in *SeMALR1* resulting in loss of function. These rearrangements did, however, not inactivate the *cis*-regulatory sequences of the *MAL* genes, since complementation with a functional *ScMAL13* allele caused induction of most *SeMAL* genes (Figure 6, Figure 7B) and, thereby, the heterotic maltotriose-positive phenotype of the hybrid strain HTSH020. Together with the high copy number of *SeAGT1*, this heterotic complementation may have been the main driver for colonization of low-temperature brewing processes by the early hybrid ancestors of current *S. pastorianus* strains. Recent work on adaptation to brewing environments of laboratory *S. cerevisiae × S. eubayanus* hybrids showed loss maltotriose utilization during serial transfer in wort (37). A similar loss of maltotriose utilization is frequently encountered in *S. cerevisiae* ale strains (54), as well as in some Saaz-type *S. pastorianus* strains (55). This is thus in contrast with retention of a maltotriose assimilation phenotype by Frohberg-type *S. pastorianus* strains. This may have been facilitated by the occurrence of multiple copies of the *SeAGT1* gene in the *S. eubayanus* ancestor, which could act as a sequence buffer to counteracting adverse effects of gene copy loss. The recent release of the first long-read sequencing assembly of *S. pastorianus* enabled a precise chromosomal mapping of the maltose-metabolism genes (56) and showed that the Frohberg type *S. pastorianus* strains CBS 1483 harbored one copy of *SeAGT1* on the *S. eubayanus* CHRXV section (as in CDFM21L.1) of the chimeric chromosome formed from SeCHRXV and SeCHRVIII (56).

Differential retention and loss of maltotriose consumption in *S. pastorianus* lineages may reflect different brewing process conditions during domestication. In modern brewing processes based on high-gravity wort, cell division is largely constrained to the glucose and maltose phases, which occur before depletion of nitrogen sources (57). It may be envisaged that, in early lager-brewing processes, unstandardized mashing processes generated wort with a higher maltotriose content, which would have allowed for continued yeast growth during the maltotriose consumption phase. During serial transfer on sugar mixtures, the selective advantage of consuming a specific sugar from a mixture correlates with the number of generations on that sugar during each cycle (58, 59). Such conditions would therefore have conferred a significant selective advantage to a maltotriose-assimilating *S. cerevisiae* x *S. eubayanus* hybrid, especially if, similar to current ale yeasts, the *S. cerevisiae* parent was unable to ferment maltotriose.

The heterotic phenotype that was reconstructed in the interspecies *S. cerevisiae* x *S. eubayanus* hybrid HTS020 resulted from combination of dominant and recessive genetic variations from both parental genomes. *S. eubayanus* contributed the *SeAGT1* gene and its functional *cis*-regulatory sequences, but also harbored recessive mutations in *MALR* genes that allowed full expression of the heterotic phenotype. These mutations were complemented with a set of *S. cerevisiae* genes including a functional *MALR* and a non-functional *ScAGT1* gene to match the mutations found in *S. pastorianus* (2). Although some *S. pastorianus* strains harbor an additional maltotriose transporter encoded by *SpMTT1* (30), this gene was recently proposed to have emerged after the original hybridization event as a result of repeated recombination between *MALT* genes from both sub-genomes (33).

Maltotriose fermentation is likely not the only heterotic phenotype of *S. pastorianus* strains. Flocculation or formation of complex aroma profiles (28, 60) are phenotypes that are not fully understood and difficult to reproduce, that also might result from heterosis (37). Laboratory-made *S. cerevisiae* x *S. eubayanus* hybrids hold great potential for brewing process intensification and for increasing product diversity. In addition to increasing our understanding of the evolutionary history of lager yeast genomes evolutionary, this study has implications for the design of new hybrids. Hitherto, laboratory crosses of *S. cerevisiae* x *S. eubayanus* strains were designed based on combination of dominant traits of the parental strains. Our results show that recessive traits can be just as important as contributors to the genetic diversity of such hybrids.

## Materials and methods

### Strains and maintenance

All strains used in this study are listed in Table 1. Stock cultures of *S. eubayanus* and *S. cerevisiae* strains were grown in YPD (10 g L^−1^ yeast extract, 20 g L^−1^ peptone and 20 g L^−1^ glucose) until late exponential phase, complemented with sterile glycerol to a final concentration of 30 % (v/v) and stored at −80 °C as 1 mL aliquots until further use.

**Table 1:** Saccharomyces strains used in this study. The abbreviation *malΔ* indicates *mal11-mal12*::loxP *mal21-mal22*::loxP *mal31-32*::loxP. Mal and Mtt denote the maltose and maltotriose phenotype respectively.

| Name | Species | Relevant genotype | Origin |
| --- | --- | --- | --- |
| CDFM21L.1 | *S. eubayanus* | Wildtype Mal<sup>-</sup> Mtt<sup>-</sup> | (38) |
| ABFM5L.1 | *S. eubayanus* | Wildtype Mal<sup>-</sup> Mtt<sup>-</sup> | (38) |
| CBS 12357 | *S. eubayanus* | Wildtype Mal<sup>+</sup> Mtt<sup>-</sup> | (1) Westerdijk institute<sup>a</sup> |
| CBS 1483 | *S. pastorianus* | Wildtype Mal<sup>+</sup> Mtt<sup>+</sup> | Westerdijk institute<sup>a</sup> |
| CEN.PK113-7D | *S. cerevisiae* | MATa MAL1x MAL2x MAL3x MAL4x MAL2-8C SUC2 LEU2 URA3 | (86) |
| IMZ616 | *S. cerevisiae* | MATa *ura3-52 LEU2 MAL2-8C malΔ mp2/3hΔ suc2Δ ima1Δ ima2Δ ima3Δ ima4Δ ima5Δ pUDC156 (Spcas9 URA3 ARS4 CEN6)* | (44) |
| IMX1365 | *S. cerevisiae* | MATa *ura3-52 LEU2 MAL2-8C malΔ mph2/3Δ suc2Δ ima1Δ ima2Δ ima3Δ ima4Δ ima5Δ pUDC156 (URA3 cas9) sga1Δ::ScTDH3<sub>p</sub>-ScMAL12-ScADH1<sub>t</sub> ScTEF1<sub>p</sub>-ScAGT1-ScCYC1<sub>t</sub>* | (13) |
| IMX1702 | *S. cerevisiae* | MATa *ura3-52 LEU2 MAL2-8C malΔ mph2/3Δ suc2Δ ima1Δ ima2Δ ima3Δ ima4Δ ima5Δ pUDC156 (URA3 cas9) sga1Δ::ScTDH3<sub>p</sub>-ScMAL12-ScADH1<sub>t</sub> ScTEF1<sub>p</sub>-SeMALT1-ScCYC1<sub>t</sub>* | This study |
| IMX1704 | *S. cerevisiae* | MATa *ura3-52 LEU2 MAL2-8C malΔ mph2/3Δ suc2Δ ima1Δ ima2Δ ima3Δ ima4Δ ima5Δ pUDC156 (URA3 cas9) sga1Δ::ScTDH3<sub>p</sub>-ScMAL12-ScADH1<sub>t</sub> ScTEF1<sub>p</sub>-SeMALT2-ScCYC1<sub>t</sub>* | This study |
| IMX1706 | *S. cerevisiae* | MATa *ura3-52 LEU2 MAL2-8C malΔ mph2/3Δ suc2Δ ima1Δ ima2Δ ima3Δ ima4Δ ima5Δ pUDC156 (URA3 cas9) sga1Δ::ScTDH3<sub>p</sub>-ScMAL12-ScADH1<sub>t</sub> ScTEF1<sub>p</sub>-SeMALT3-ScCYC1<sub>t</sub>* | This study |
| IMX 1708 | *S. cerevisiae* | MATa *ura3-52 LEU2 MAL2-8C malΔ mph2/3Δ suc2Δ ima1Δ ima2Δ ima3Δ ima4Δ ima5Δ pUDC156 (URA3 cas9) sga1Δ::ScTDH3<sub>p</sub>-ScMAL12-ScADH1<sub>t</sub> ScTEF1<sub>p</sub>-SeAGT1-ScCYC1<sub>t</sub>* | This study |
| IMX1313 | *S. cerevisiae* | MATa *ura3-52 LEU2 MAL2-8C malΔ mph2/3Δ suc2Δ ima1Δ ima2Δ ima3Δ ima4Δ ima5Δ sga1Δ::ScTEF1<sub>p</sub>-ScMAL31-ScCYC1<sub>t</sub> pUDC156 (URA3 cas9)* | This study |
| IMX1313Δ | *S. cerevisiae* | MATa *ura3-52 LEU2 MAL2-8C malΔ mph2/3Δ suc2Δ ima1Δ ima2Δ ima3Δ ima4Δ ima5Δ sga1Δ::ScTEF1<sub>p</sub>-ScMAL31-ScCYC1<sub>t</sub>* | This study |
| IMZ752 | *S. cerevisiae* | MATa *ura3-52 LEU2 MAL2-8C malΔ mph2/3Δ suc2Δ ima1Δ ima2Δ ima3Δ ima4Δ ima5Δ sga1Δ::ScTEF1<sub>p</sub>-ScMAL31-ScCYC1<sub>t</sub> pUDE843 (ori (ColE1) bla 2μ ScTDH3<sub>p</sub>-SeMALS1-ScADH1<sub>t</sub> URA3)* | This study |
| IMZ753 | *S. cerevisiae* | MATa *ura3-52 LEU2 MAL2-8C malΔ mph2/3Δ suc2Δ ima1Δ ima2Δ ima3Δ ima4Δ ima5Δ sga1Δ::ScTEF1<sub>p</sub>-ScMAL31-ScCYC1<sub>t</sub> pUDE844 (ori (ColE1) bla 2μ ScTDH3<sub>p</sub>-SeMALS2-ScADH1<sub>t</sub> URA3)* | This study |
| IMK820 | *S. eubayanus* | MATa/MATα *Semalt1Δ/Semalt1Δ* | This study |
| IMK823 | *S. eubayanus* | MATa/MATα *Seagt1Δ/Seagt1Δ (X3)* | This study |
| IMX1939 | *S. eubayanus* | MATa/MATα *Semalt1Δ/Semalt1Δ Sesga1Δ::ScMAL13/Sesga1Δ::ScMAL13* | This study |
| IMX1940 | *S. eubayanus* | MATa/MATα *Seagt1Δ/Seagt1Δ Sesga1Δ::ScMAL13/Sesga1Δ::ScMAL13* | This study |
| IMX1762 | *S. eubayanus* | MATa/MATα *Sesga1Δ::ScMAL12/Sesga1Δ::ScMAL12* | This study |
| IMX1765 | *S. eubayanus* | MATa/MATα *Sesga1Δ::ScMAL13/Sesga1Δ::ScMAL13* | This study |
| CBC-1 | *S. cerevisiae* | MATa/MATα Mal<sup>+</sup> Mtt<sup>-</sup> | Lallemand |
| HTSH020 | *S. cerevisiae* X *S. eubayanus* | MATa/MATα Mal<sup>+</sup> Mtt<sup>+</sup> | This study |

<sup>a</sup> Westerdijk Fungal Biodiversity Institute www.westerdijk.nl/

### Media and cultivation

*S. eubayanus* batch cultures were grown on synthetic medium (SM) containing 3.0 g L^−1^ KH_2_PO_4_, 5.0 g L^−1^ (NH_4_)_2_SO_4_, 0.5 g L^−1^ MgSO_4_.7H_2_O, 1 mL L^−1^ trace element solution, and 1 mL L^−1^ vitamin solution (61). The pH was set to 6.0 with 2 M KOH prior to autoclaving at 120 °C for 20 min. Vitamin solutions were sterilized by filtration and added to the sterile medium. Concentrated sugar solutions were autoclaved at 110 °C for 20 min or filter sterilized and added to the sterile flasks to give a final concentration of 20 g L^−1^ glucose (SMG), maltose (SMM) or maltotriose (SMMt). With the exception of IMZ752 and IMZ753, *S. cerevisiae* batch cultures were grown on SM supplemented with 150 mg L^−1^ uracil (62) to compensate for loss of plasmid pUDC156 that carried the *Spcas9* endonuclease gene, and supplemented with 20 g L^−1^ glucose (SM_u_G), maltose (SM_u_M) or maltotriose (SM_u_Mt). All batch cultures were grown in 250 mL shake flasks with a working volume of 50 mL. The cultures were inoculated at an initial OD_660nm_ of 0.1 and incubated under an air atmosphere and shaken at 200 rpm and at 20 °C in a New Brunswick™ Innova44 incubator (Eppendorf Nederland B.V, Nijmegen, The Netherlands).

*S. eubayanus* strains transformed with plasmids pUDP052 (gRNA_*SeSGA1*_), pUDP091 (gRNA_*SeMALT1*_) and pUDP090 (gRNA_*SeAGT1*_) were selected on modified SMG medium (SM_Ace_G) in which (NH_4_)_2_SO_4_ was replaced by 6.6 g L^−1^ K_2_SO_4_ and 10 mM acetamide (63). SM-based solid media contained 2 % Bacto Agar (BD Biosciences, Franklin Lakes, NJ). *S. cerevisiae* strains expressing either *SeMALT, SeMALS* or *ScMALR* were selected on SM_Ace_G. For plasmid propagation, *E. coli* XL1-Blue-derived strains (Agilent Technologies, Santa Clara, CA) were grown in Lysogeny Broth medium (LB, 10 g L^−1^ tryptone, 5 g L^−1^ yeast extract, 5 g L^−1^ NaCl) supplied with 100 mg L^−1^ ampicillin. Synthetic wort medium (SWM) for growth studies contained 14.4 g·L^−1^ glucose, 2.3 g·L^−1^ fructose, 85.9 g·L^−1^ maltose, 26.8 g·L^−1^ maltotriose, 5 g·L^−1^ (NH_4_)_2_SO_4_, 3 g·L^−1^ KH_2_PO_4_, 0.5 g·L^−1^ MgSO_4_.7H_2_O, 1 mL·L^−1^ trace element solution, 1 mL·L^−1^ vitamin solution, supplemented with the anaerobic growth factors ergosterol and Tween 80 (0.01 g·L^−1^ and 0.42 g·L^−1^ respectively), as previously described (61).

Industrial wort (containing 14.4 g L^−1^ glucose, 85.9 g L^−1^ maltose, 26.8 g L^−1^ maltotriose, 2.3 g L^−1^ fructose and 269 mg L^−1^ FAN) was provided by Heineken Supply Chain B.V. (Zoeterwoude, the Netherlands). The wort was supplemented with 1.5 mg L^−1^ of Zn^2+^ by addition of ZnSO_4_.7H_2_O, autoclaved for 30 min at 121 □C, filtered using Nalgene 0.2 μm SFCA bottle top filters (Thermo Scientific) and diluted with sterile demineralized water. Sporulation medium consisted of 2 % (w/v) KAc in MilliQ water set to pH 7.0 with KOH, autoclaved at 121 °C for 20 min.

### Microaerobic growth experiments

Microaerobic cultures were grown in 250-mL airlock-capped Neubor infusion bottles (38 mm neck, Dijkstra, Lelystad, Netherlands) containing 200 mL three-fold diluted industrial wort supplemented with 0.4 mL L^−1^ Pluronic antifoam (Sigma-Aldrich, St. Louis, MO). Bottle caps were equipped with a 0.5 mm x 16 mm Microlance needle (BD Biosciences) sealed with cotton to prevent pressure build-up. Sampling was performed aseptically with 3.5 mL syringes using a 0.8 mm x 50 mm Microlance needle (BD Biosciences). Microaerobic cultures were inoculated at an OD_660nm_ of 0.1 from stationary-phase precultures in 50 mL Bio-One Cellstar Cellreactor tubes (Sigma-Aldrich) containing 30 mL of the same medium, grown for 4 days at 12 °C. Bottles were incubated at 12 °C and shaken at 200 rpm in a New Brunswick Innova43/43R shaker (Eppendorf Nederland B.V.). At regular intervals, 3.5 mL samples were collected in 24 deep-well plates (EnzyScreen BV, Heemstede, Netherlands) using a LiHa liquid handler (Tecan, Männedorf, Switzerland) to measure OD_660nm_ and external metabolites. 30 μL of each sample was diluted 5-fold in demineralized water in a 96 well plate and OD_660nm_ was measured with a Magellan Infinite 200 PRO spectrophotometer (Tecan). From the remaining sample, 150 μL was vacuum filter sterilized using 0.2 μm Multiscreen filter plates (Merck, Darmstadt, Germany) for HPLC measurements.

### Analytical methods

Optical densities of yeast cultures were measured with a Libra S11 spectrophotometer (Biochrom, Cambridge, United Kingdom) at a wavelength of 660 nm. Biomass dry weight was measured by filtering 10-mL culture samples over pre-weighed nitrocellulose filters with a pore size of 0.45 μm. Filters were washed with 10 mL water, dried in a microwave oven (20 min at 350 W) and reweighed. Sugars were measured using a high pressure liquid chromatography Agilent Infinity 1260 series (Agilent Technologies) using a Bio-Rad Aminex HPX-87H column at 65 °C with 5 mM sulfuric acid at a flow rate of 0.8 mL min^−1^. Compounds were measured using a RID at 35 °C. Samples were centrifuged at 13,000 g for 5 min to collect supernatant or 0.2 μm filter-sterilized before analysis.

### Plasmid construction

Plasmids used and constructed in this study are listed in Table 2, oligonucleotide primers used in this study are listed in Supplementary Table 1. Coding regions of *SeMALT1, SeMALT2, SeMALT3* and *SeAGT1* were amplified from CDFM21L.1 genomic DNA with Phusion High-Fidelity DNA polymerase (Thermo Scientific), according to the supplier’s instructions with primers pairs 12355/12356, 12357/12358, 12359/12360 and 12361/12362, respectively. The coding sequence of *ScMAL31* was amplified from CEN.PK113-7D genomic DNA with Phusion High-Fidelity DNA polymerase (Thermo Scientific), according to the supplier’s instructions with primer pairs 9942/9943. Each primer carried a 40 bp extension complementary to the plasmid backbone of p426-TEF-amdS (64), which was PCR amplified using Phusion High-Fidelity DNA polymerase (Thermo Scientific) and primers 7812/5921. Each transporter fragment was assembled with the p426-TEF-amdS backbone fragment using NEBuilder HiFi DNA Assembly (New England Biolabs, Ipswich, MA), resulting in plasmids pUD444 (*ScMAL31*), pUD794 (*SeMALT1*), pUD795 (*SeMALT2*), pUD796 (*SeMALT3*) and pUD797 (*SeAGT1*). All plasmids were verified for correct assembly by Sanger sequencing (Baseclear, Leiden, The Netherlands).

**Table 2.**
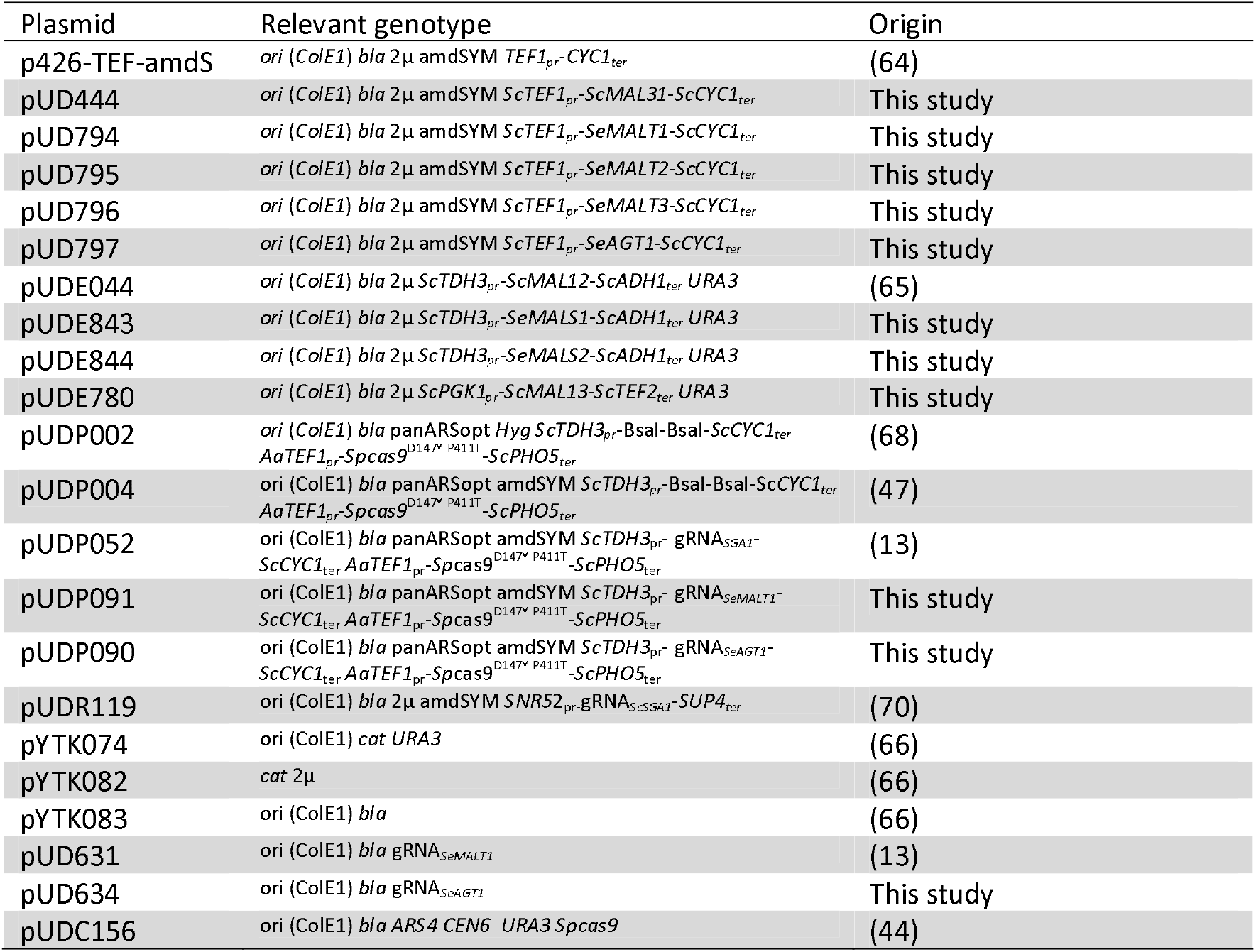
Plasmids used in this study

*SeMALS1* and *SeMALS2* were amplified from CDFM21L.1 genomic DNA with Phusion High-Fidelity DNA polymerase (Thermo Scientific), with primers pairs 14451/14453 and 14452/14453, respectively. Each primer pair carried a 30 bp extension complimentary to the plasmid backbone of pUDEO44 (65) which was PCR amplified using Phusion High-Fidelity DNA polymerase (Thermo Scientific) and primers 14449/14450. Resulting amplicons were assembled using NEBuilder HiFi DNA Assembly (New England Biolabs), resulting in plasmids pUDE843 (*SeMALS1*) and pUDE844 (*SeMALS2*) that were verified by Sanger sequencing (Baseclear).

*S. cerevisiae ScMAL13*, the *ScPGK1* promoter and the *ScTEF2* terminator were amplified from CEN.PK113-7D genomic DNA with Phusion High-Fidelity DNA polymerase (Thermo Scientific), with primer pairs 12915/12916, 9421/9422 and 10884/10885, respectively. Fragments were gel purified and used with pYTK074, pYTK082 and pYTK083 in Golden Gate assembly according to the yeast toolkit protocol (66) resulting in pUDE780, which was verified by Sanger sequencing (Baseclear).

Guide-RNA (gRNA) sequences for deletion of *SeMALT1* and *SeAGT1* in CDFM21L.1 were designed as described previously (47). The DNA sequences encoding these gRNAs were synthesized at GeneArt (Thermo Scientific) and were delivered in pUD631 and pUD634, respectively. The gRNA spacer sequences (*SeMALT1* 5’ CCCCGATATTCTTTACACTA 3’, *SeAGT1* 5’-AGCTTTGCGAAAATATCCAA-3’) and the structural gRNA sequence were flanked at their 5’ ends by the Hammerhead ribozyme (HH) and at their 3’ ends by the Hepatitis Delta Virus ribozyme (HDV) (67). The HH-gRNA-HDV fragment was flanked on both ends with a BsaI site for further cloning (47, 68). Plasmids pUDP091 (gRNA_*SeMALT1*_) and pUDP090 (gRNA_*SeAGT1*_) were constructed by Golden Gate cloning by digesting pUDP004 and the gRNA-carrying plasmid (pUD631 and pUD634, respectively) using BsaI and ligating with T4 ligase (69). Correct assembly was verified by restriction analysis with PdmI (Thermo Scientific) and Sanger sequencing (Baseclear).

### Strain construction

*S. cerevisiae* IMZ616, which cannot grow on α-glucosides (44), was used as a host to test functionality of individual *S. eubayanus* (putative) maltose transporter genes (13). *S. cerevisiae* IMX1702 was constructed by integrating *ScTDH3_p_-ScMAL12-ScADH1_t_* and *ScTEF1_p_-SeMALT1-ScCYC1_t_* at the *ScSGA1* locus of strain IMZ616. A fragment containing the *ScTDH3_p_-ScMAL12-ScADH1_t_* transcriptional unit was PCR amplified using Phusion High-Fidelity DNA polymerase (Thermo Scientific) from pUDE044 with primers 9596/9355, which included a 5’ extension homologous to the upstream region of the *SeSGA1* locus and an extension homologous to the co-transformed transporter fragment, respectively. The DNA fragment carrying the *S. eubayanus SeMALT1* maltose symporter (*ScTEF1_p_-SeMALT1-ScCYC1_t_*) was PCR amplified from pUD794 using primers 9036/9039, which included a 5’ extension homologous to the co-transformed transporter fragment and an extension homologous to the downstream region of the *ScSGA1* locus, respectively. To facilitate integration in strain IMZ616, the two PCR fragments were co-transformed with plasmid pUDR119 (*amdS*), which expressed a gRNA targeting *ScSGA1* (spacer sequence: 5’-ATTGACCACTGGAATTCTTC-3’) (70). The plasmid and repair fragments were transformed using the LiAc yeast transformation protocol (71) and transformed cells were plated on SM_Ace_G. Correct integration was verified by diagnostic PCR with primers pairs 4226/4224. Strains *S. cerevisiae* IMX1704, IMX1706 and IMX1708 were constructed following the same principle, but instead of using pUD794 to generate the transporter fragment, pUD795, pUD796 and pUD797 were used to PCR amplify *ScTEF1_p_-SeMALT2-ScCYC1_t_, ScTEF1_p_-SeMALT3-ScCYC1_t_* and *ScTEF1_p_-SeAGT1-ScCYC1_t_* respectively. IMX1313 was constructed in a similar way using only *ScTEF1_p_-ScMAL31-ScCYC1_t_* amplified with primer pair 9036/11018 which contain 5’and 3’ extensions homologous to the upstream and downstream region of the *ScSGA1* locus. Correct integration was verified by diagnostic PCR with primer pair 4226/4224 (Supplementary Figure 1). All PCR-amplified gene sequences were Sanger sequenced (BaseClear). IMX1313 was grown on YPD to loose pUDR119 (*URA3*) and pUDC156 (*amdS*). An isolate unable to grown on SMG without uracil and with acetamide was selected and named IMX1313Δ. This strain was able to grow on SMG supplemented with 150 mg L^−1^ uracil.

To assess functionality of CDFM21L.1 *SeMALS1*, IMX1313Δ was transformed with 100 ng pUDE843 (*ScTDH3_p_-SeMALS1-ScADH1_t_*) by electroporation (47), resulting in strain IMZ752. Transformants were selected on SMG plates after 5 days of incubation at 20 °C and validated by PCR (DreamTaq polymerase, Thermo Scientific) using primer pair 14454/14455 (Supplementary Figure 1). Similarly, functionality of the *SeMALS2* maltase gene of CDFM21L.1 was assessed by transforming IMX1313Δ with pUDE844 (*ScTDH3*_pr_-*SeMALS2-ScADH1_ter_*), resulting in strain IMZ753.

*S. eubayanus* IMK820 (*SemalT1*Δ) was constructed by transforming CDFM21L.1 with 200 ng of pUDP091 and 1 μg of 120 bp repair fragment obtained by mixing an equimolar amount of primers 12442/12443, as previously described (47). As a control, the same transformation was performed without including the repair DNA fragment. Transformants were selected on SM_Ace_G plates. *S. eubayanus* IMK823 (*Seagt1*Δ) was constructed similarly, using pUDP090 and primer pair 11320/11321. Deletion of *SemalT1* was verified by PCR with primer pair 11671/11672 and Sanger sequencing. The *Seagt1* deletion was verified by PCR using primer pair 12273/12274, and by Illumina whole-genome sequencing and read alignment to the reference genome of CDFM21L.1 (Bioproject accession number PRJNA528469).

Strains IMX1765, IMX1939 and IMX1940 were constructed by inserting *ScPGK1_p_-ScMAL13-ScTEF2_t_* at the *SeSGA1* locus of CDFM21L.1, IMK820 and IMK823, respectively. A repair fragment containing *ScPGK1_p_-ScMAL13-ScTEF2_t_* was amplified from pUDE780 with primer pair 12917/12918. Strains CDFML21L.1, IMK820 and IMK823 were transformed by electroporation by addition of 350 ng of repair fragment and 560 ng pUDP052 (*amdS*) into the cells as previously described (47). Transformants were plated on SM_Ace_G and incubated at 20 °C. IMX1762 was constructed similarly using a repair fragment with *ScTDH3_p_-ScMAL12-ScADH1_t_* amplified from pUDE044 with primer pair 12319/12320. Strains were verified by PCR using primer pair 12635/12636 and Sanger sequencing.

### Hybrid Construction

The *S. cerevisiae* X *S. eubayanus* hybrid HTSH020 was constructed by spore-to-spore mating. The *S. eubayanus* strain CDFM21L.1 and the *S. cerevisiae* strain CBC-1 were grown in 20 mL YPD at 20 °C until late exponential phase. Cells were centrifuged for 5 min at 1000 g and washed twice in demineralized water. Cells were re-suspended in 20 mL sporulation medium and incubated for 64 h at 20 °C. Presence of spores was verified by microscopy. Asci were harvested by centrifugation for 5 min at 1000 g, and washed with demineralized water, resuspended in 100 μL demineralized water containing 100 U/mL of Zymolyase (MP Bio, Santa Ana, CA) and incubated for 10 min at 30 °C. Spores were washed and plated on the edge of a YPD agar plate. Spores from the two strains were brought in contact with each other with an MSM System 400 micromanipulator (Singer Instruments, Watchet, United Kingdom). Zygote formation was observed after 6-8 h. Emerging colonies were re-streaked twice on SM 2% maltose at 12 °C. Successful hybridization was verified by multiplex PCR using DreamTaq DNA polymerase (Thermo Scientific), by amplifying the *S. cerevisiae* specific *MEX67* gene with primer pairs 8570/8571 and by amplifying the *S. eubayanus* specific gene *SeFSY1* with primers 8572/8573 (Supplementary Figure 2), as previously described (72).

### Illumina sequencing

Genomic DNA of *S. eubayanus* strains CDFM21L.1 and ABFM5L.1, *S. cerevisiae* strain CBC-1 and *S. cerevisiae* x *S. eubayanus* strain HTSH020 was isolated as previously described (4). Paired-end sequencing (2 × 150 bp) was performed on a 350 bp PCR-free insert library using Illumina HiSeq2500 (San Diego, CA) by Novogene(HK) Company Ltd (Hong Kong, China). Genomic DNA of the strains CBC-1 and HTSH020 was sequenced in house on a MiSeq sequencer (Illumina) with 300 bp paired-end reads using PCR-free library preparation. Sequence data are available at NCBI under Bioproject accession number PRJNA528469.

### MinION long read sequencing

For long-read sequencing, a 1D sequencing library (SQK-LSK108) was prepared for CDFM21L.1 and CBC-1 and loaded onto an FLO-MIN106 (R9.4) flow cell, connected to a MinION Mk1B unit (Oxford Nanopore Technology, Oxford, United Kingdom), according to the manufacturer’s instructions. MinKNOW software (version 1.5.12; Oxford Nanopore Technology) was used for quality control of active pores and for sequencing. Raw files generated by MinKNOW were base-called using Albacore (version 1.1.0; Oxford Nanopore Technology). Reads with a minimum length of 1000 bp were extracted in fastq format. For CDFM21L.1, 3.26 Gb of sequence with an average read length of 8.07 kb was obtained and for CBC-1 3.04 Gb sequence with an average read length of 7.27 kb. Sequencing data are available at NCBI under Bioproject accession number PRJNA528469.

### De novo *assembly*

*De novo* assembly of the Oxford Nanopore MinION dataset was performed using Canu (v1.4, setting: genomesize=12m) (73). Assembly correctness was assessed using Pilon (74) and further corrected by “polishing” of sequencing/assembly errors by aligning Illumina reads with BWA (75) using correction of only SNPs and short indels (–fix bases parameter). For HTSH020, an artificial reference genome was made by combining the assembly of CBC-1 and CDFM21L.1. The genome assemblies were annotated using the MAKER2 annotation pipeline (version 2.31.9) (76), using SNAP (version 2013-11-29) (77) and Augustus (version 3.2.3) (78) as *ab initio* gene predictors. *S. cerevisiae* S288C EST and protein sequences were obtained from SGD (*Saccharomyces* Genome Database, http://www.yeastgenome.org/) and were aligned using BLASTX on the obtained polished sequence assembly (BLAST version 2.2.28+) (79). Predicted translated protein sequences of the final gene model were aligned to the *S. cerevisiae* S288C protein Swiss-Prot database using BLASTP (http://www.uniprot.org/). Custom-made Perl scripts were used to map systematic names to the annotated gene names (Supplementary Table 2). Error rates in nanopore-sequencing data were estimated from the q score (Phred scaled) per read, as calculated by the base caller Albacore (version 1.1.0) (Oxford Nanopore Technology). Average q score was used to calculate the error P= 10^q/10^.

### RNA isolation

CDFM21L1, IMX1765, IMX1939 and HTS020 were grown in SMG, SMM and SMMt until mid-exponential phase (OD_660nm_ of 12 for SMG/SMM and of OD_660nm_ 15 for SMMt). Culture samples corresponding to ca. 200 mg of biomass dry weight were directly quenched in liquid nitrogen. The samples were processed and total RNA extracted as previously described (80). Prior to cDNA synthesis, purity, concentration and integrity of the RNA in the samples was assessed with Nanodrop (Thermo Scientific), Qubit (Thermo Scientific) and Tapestation 220 with RNA Screen Tape (Agilent Technologies), respectively, according the manufacturers’ recommendations. cDNA libraries were prepared using the TruSeq RNA V2 kit (Illumina). Paired-end sequencing (2 × 150 bp) was performed on a 300 bp PCR-free insert library on a HISeq 2500 (Illumina) at Novogene (HK) Company Ltd (Hong Kong, China). Duplicate biological samples were processed, generating an average sequence quantity of 23.7M reads per sample. Reads were aligned to the CDFM21L.1 reference assembly (GEO (https://www.ncbi.nlm.nih.gov/geo/) accession number GSE133146) using a two-pass STAR (81) procedure. In the first pass, splice junctions were assembled and used to inform the second round of alignments. Introns between 15 and 4000 bp were allowed, and soft clipping was disabled to prevent low-quality reads from being spuriously aligned. Ambiguously-mapped reads were removed from the dataset. Expression level for each transcript were quantified using htseq-count (82) in union mode. Fragments per kilo-base of feature (gene) per million reads mapped (FPKM) values were calculated by “Applying the fpkm method” from the edgeR package (83, 84). Differential expression analysis was performed using DESeq (85).

### Accession Numbers

The sequencing data were deposited at NCBI (https://www.ncbi.nlm.nih.gov/) under the Bioproject PRJNA528469 and the transcriptomics data were deposited at GEO (Genome Expression Omnibus https://www.ncbi.nlm.nih.gov/geo/) under accession number GSE133146.

## Supporting information

Supplemntary figures with captions

## Supplementary Data

Supplementary Data are available

## Authors contributions

JMD conceived the study and designed the experiments. NB, AB, LvbE, ARGdV, SMW and JD performed the experimental work. NB and MvdB performed bioinformatics analysis. NB, AB, FYB, JTP and JMGD supervised the study and wrote the manuscript. All authors read and approved the final manuscript.

## Acknowledgements

We thank Flip de Groot for his technical assistance in gene editing in *S. eubayanus*, Niels G.A. Kuijpers, Viktor M. Boer and Jan-Maarten Geertman for their critical input.

## References

1. Libkind D, Hittinger CT, Valério E, Gonçalves C, Dover J, Johnston M, Gonsalves P, Sampaio JP. 2011. Microbe domestication and the identification of the wild genetic stock of lager-brewing yeast. Proc Natl Acad Sci U S A:201105430.

2. Nakao Y, Kanamori T, Itoh T, Kodama Y, Rainieri S, Nakamura N, Shimonaga T, Hattori M, Ashikari T. 2009. Genome sequence of the lager brewing yeast, an interspecies hybrid. DNA Res 16:115–129.

3. Walther A, Hesselbart A, Wendland J. 2014. Genome sequence of *Saccharomyces carlsbergensis*, the world’s first pure culture lager yeast. G3 (Bethesda) 4:783–793.

4. Van den Broek M, Bolat I, Nijkamp JF, Ramos E, Luttik MAH, Koopman F, Geertman JM, De Ridder D, Pronk JT, Daran J-MG. 2015. Chromosomal copy number variation in *Saccharomyces pastorianus* evidence for extensive genome dynamics in industrial lager brewing strains. Appl Environ Microbiol:AEM. 01263–01215.

5. Krogerus K, Magalhães F, Vidgren V, Gibson B. 2015. New lager yeast strains generated by interspecific hybridization. J Ind Microbiol Biotechnol 42:769–778.

6. Hebly M, Brickwedde A, Bolat I, Driessen MR, de Hulster EA, van den Broek M, Pronk JT, Geertman J-M, Daran J-MG, Daran-Lapujade P. 2015. S. *cerevisiae* ϗ S. *eubayanus* interspecific hybrid, the best of both worlds and beyond. FEMS Yeast Res 15.

7. Krogerus K, Magalhães F, Vidgren V, Gibson B. 2017. Novel brewing yeast hybrids: creation and application. Appl Environ Microbiol 101:65–78.

8. Zastrow C, Hollatz C, De Araujo P, Stambuk B. 2001. Maltotriose fermentation by *Saccharomyces cerevisiae*. J Ind Microbiol Biotechnol 27:34–38.

9. Alves SL, Herberts RA, Hollatz C, Trichez D, Miletti LC, De Araujo PS, Stambuk BU. 2008. Molecular analysis of maltotriose active transport and fermentation by *Saccharomyces cerevisiae* reveals a determinant role for the AGT1 permease. Appl Environ Microbiol 74:1494–1501.

10. Vidgren V, Huuskonen A, Virtanen H, Ruohonen L, Londesborough J. 2009. Improved fermentation performance of a lager yeast after repair of its *AGT1* maltose and maltotriose transporter genes. Appl Environ Microbiol 75:2333–2345.

11. Brickwedde A, van den Broek M, Geertman J-MA, Magalhães F, Kuijpers NG, Gibson B, Pronk JT, Daran J-MG. 2017. Evolutionary engineering in chemostat cultures for improved maltotriose fermentation kinetics in *Saccharomyces pastorianus* lager brewing yeast. Front Microbiol 8:1690.

12. Zheng X, D’Amore T, Russell I, Stewart G. 1994. Factors influencing maltotriose utilization during brewery wort fermentations. J Am Soc Brew Chem 52:41–47.

13. Brickwedde A, Brouwers N, van den Broek M, Gallego Murillo JS, Fraiture JL, Pronk JT, Daran J-MG. 2018. Structural, physiological and regulatory analysis of maltose transporter genes in *Saccharomyces eubayanus* CBS 12357T. Front Microbiol 9:1786.

14. Baker EP, Hittinger CT. 2018. Evolution of a novel chimeric maltotriose transporter in *Saccharomyces eubayanus* from parent proteins unable to perform this function. bioRxiv doi: 10.1101/431171.

15. Krogerus K, Magalhães F, Vidgren V, Gibson B. 2017. Novel brewing yeast hybrids: creation and application. Appl Microbiol Biotechnol 101:65–78.

16. Krogerus K, Magalhães F, Vidgren V, Gibson B. 2015. New lager yeast strains generated by interspecific hybridization. Journal of industrial microbiology & biotechnology 42:769–778.

17. Nikulin J, Krogerus K, Gibson B. 2018. Alternative Saccharomyces interspecies hybrid combinations and their potential for low-temperature wort fermentation. Yeast (Chichester, England) 35:113–127.

18. Cheng Q, Michels CA. 1991. *MAL11* and *MAL61* encode the inducible high-affinity maltose transporter of *Saccharomyces cerevisiae*. J Bacteriol 173:1817–1820.

19. Vidgren V, Multanen JP, Ruohonen L, Londesborough J. 2010. The temperature dependence of maltose transport in ale and lager strains of brewer’s yeast. FEMS Yeast Res 10:402–411.

20. Naumov GI, Naumova ES, Michels C. 1994. Genetic variation of the repeated *MAL* loci in natural populations of *Saccharomyces cerevisiae* and *Saccharomyces paradoxus*. Genetics 136:803–812.

21. Charron MJ, Read E, Haut SR, Michels CA. 1989. Molecular evolution of the telomere-associated *MAL* loci of *Saccharomyces*. Genetics 122:307–316.

22. Hayford A, Jespersen L. 1999. Characterization of Saccharomyces cerevisiae strains from spontaneously fermented maize dough by profiles of assimilation, chromosome polymorphism, PCR and MAL genotyping. Journal of applied microbiology 86:284–294.

23. Bell PJ, Higgins VJ, Dawes IW, Bissinger PH. 1997. Tandemly repeated 147 bp elements cause structural and functional variation in divergent MAL promoters of Saccharomyces cerevisiae. Yeast 13:1135–1144.

24. Barnett JA. 1976. The Utilization of Sugars by Yeasts1, p 125–234, Advances in carbohydrate chemistry and biochemistry, vol 32. Elsevier.

25. Salazar AN, Gorter de Vries AR, van den Broek M, Wijsman M, de la Torre Cortés P, Brickwedde A, Brouwers N, Daran J-MG, Abeel T. 2017. Nanopore sequencing enables near-complete de novo assembly of *Saccharomyces cerevisiae* reference strain CEN. PK113-7D. FEMS Yeast Res 17.

26. Baker E, Wang B, Bellora N, Peris D, Hulfachor AB, Koshalek JA, Adams M, Libkind D, Hittinger CT. 2015. The genome sequence of *Saccharomyces eubayanus* and the domestication of lager-brewing yeasts. Mol Biol Evol 32:2818–2831.

27. Han EK, Cotty F, Sottas C, Jiang H, Michels CA. 1995. of *AGT1* encoding a general a-glucoside transporter from *Saccharomyces*. Mol Microbiol 17:1093–1107.

28. Vidgren V, Gibson B. 2018. Trans-regulation and localization of orthologous maltose transporters in the interspecies lager yeast hybrid. FEMS yeast research 18:foy065.

29. Salema-Oom M, Pinto VV, Gonçalves P, Spencer-Martins I. 2005. Maltotriose utilization by industrial *Saccharomyces strains:* characterization of a new member of the a-glucoside transporter family. Appl Environ Microbiol 71:5044–5049.

30. Dietvorst J, Londesborough J, Steensma H. 2005. Maltotriose utilization in lager yeast strains: *MTT1* encodes a maltotriose transporter. Yeast 22:775–788.

31. Cousseau F, Alves Jr S, Trichez D, Stambuk B. 2013. Characterization of maltotriose transporters from the *Saccharomyces eubayanus* subgenome of the hybrid *Saccharomyces pastorianus* lager brewing yeast strain Weihenstephan 34/70. Lett Appl Microbiol 56:21–29.

32. Nguyen H-V, Legras J-L, Neuvéglise C, Gaillardin C. 2011. Deciphering the hybridisation history leading to the lager lineage based on the mosaic genomes of *Saccharomyces bayanus* strains NBRC1948 and CBS380T. PLoS One 6:e25821.

33. Brouwers N, Gorter de Vries AR, van den Broek M, Weening SM, Elink Schuurman TD, Kuijpers NGA, Pronk JT, Daran J-MG. 2018. In vivo recombination of *Saccharomyces eubayanus* maltose-transporter genes yields a chimeric transporter that enables maltotriose fermentation. bioRxiv doi:10.1101/428839.

34. Brouwers N, Gorter de Vries AR, van den Broek M, Weening SM, Elink Schuurman TD, Kuijpers NGA, Pronk JT, Daran JG. 2019. In vivo recombination of Saccharomyces eubayanus maltose-transporter genes yields a chimeric transporter that enables maltotriose fermentation. PLoS Genet 15:e1007853.

35. Baker EP, Hittinger CT. 2019. Evolution of a novel chimeric maltotriose transporter in Saccharomyces eubayanus from parent proteins unable to perform this function. PLoS Genet 15:e1007786.

36. Mertens S, Steensels J, Saels V, De Rouck G, Aerts G, Verstrepen KJ. 2015. A large set of newly created interspecific yeast hybrids increases aromatic diversity in lager beers. Appl Environ Microbiol:AEM. 02464–02415.

37. Gorter de Vries A, Voskamp MA, van Aalst ACA, Kristensen LH, Jansen L, van den Broek M, Salazar AN, Brouwers N, Abeel T, Pronk JT, Daran J-MG. 2018. Laboratory evolution of a Saccharomyces cerevisiae x S. eubayanus hybrid under simulated lager-brewing conditions: genetic diversity and phenotypic convergence. bioRxiv.

38. Bing J, Han P-J, Liu W-Q, Wang Q-M, Bai F-Y. 2014. Evidence for a Far East Asian origin of lager beer yeast. Curr Biol 24:R380–R381.

39. Peris D, Langdon QK, Moriarty RV, Sylvester K, Bontrager M, Charron G, Leducq J-B, Landry C, Libkind D, Hittinger CT. 2016. Complex ancestries of lager-brewing hybrids were shaped by standing variation in the wild yeast Saccharomyces eubayanus. PLoS Genet 12:20.

40. Darling ACE, Mau B, Blattner FR, Perna NT. 2004. Mauve: multiple alignment of conserved genomic sequence with rearrangements. Genome research 14:1394–1403.

41. Brown CA, Murray AW, Verstrepen KJ. 2010. Rapid expansion and functional divergence of subtelomeric gene families in yeasts. Curr Biol 20:895–903.

42. Moller M, Habig M, Freitag M, Stukenbrock EH. 2018. Extraordinary Genome Instability and Widespread Chromosome Rearrangements During Vegetative Growth. Genetics 210:517–529.

43. Gordon JL, Byrne KP, Wolfe KH. 2009. Additions, losses, and rearrangements on the evolutionary route from a reconstructed ancestor to the modern *Saccharomyces cerevisiae* genome. PLoS Genet 5:e1000485.

44. Marques WL, Mans R, Henderson RK, Marella ER, ter Horst J, de Hulster E, Poolman B, Daran J-M, Pronk JT, Gombert AK. 2018. Combined engineering of disaccharide transport and phosphorolysis for enhanced ATP yield from sucrose fermentation in Saccharomyces cerevisiae. Metabolic engineering 45:121–133.

45. Meurer M, Chevyreva V, Cerulus B, Knop M. 2017. The regulatable MAL32 promoter in Saccharomyces cerevisiae: characteristics and tools to facilitate its use. Yeast 34:39–49.

46. Levine J, Tanouye L, Michels CA. 1992. The UAS(MAL) is a bidirectional promotor element required for the expression of both the MAL61 and MAL62 genes of the Saccharomyces MAL6 locus. Curr Genet 22:181–189.

47. Gorter de Vries AR, Groot PA, Broek M, Daran J-MG. 2017. CRISPR-Cas9 mediated gene deletions in lager yeast *Saccharomyces pastorianus*. Microb Cell Fact 16:222.

48. Vidgren V, Kankainen M, Londesborough J, Ruohonen L. 2011. Identification of regulatory elements in the AGT1 promoter of ale and lager strains of brewer’s yeast. Yeast 28:579–594.

49. Dunn B, Sherlock G. 2008. Reconstruction of the genome origins and evolution of the hybrid lager yeast Saccharomyces pastorianus. Genome Res 18:1610–1623.

50. Okuno M, Kajitani R, Ryusui R, Morimoto H, Kodama Y, Itoh T. 2016. Next-generation sequencing analysis of lager brewing yeast strains reveals the evolutionary history of interspecies hybridization. DNA Res 23:67–80.

51. Monerawela C, Bond U. 2017. Brewing up a storm: The genomes of lager yeasts and how they evolved. Biotechnology Advances 35:512–519.

52. Alsammar HF, Naseeb S, Brancia LB, Gilman RT, Wang P, Delneri D. 2019. Targeted metagenomics approach to capture the biodiversity of Saccharomyces genus in wild environments. Environmental Microbiology Reports 11:206–214.

53. Vidgren V, Ruohonen L, Londesborough J. 2005. Characterization and functional analysis of the MAL and MPH Loci for maltose utilization in some ale and lager yeast strains. Appl Environ Microbiol 71:7846–7857.

54. Gallone B, Steensels J, Prahl T, Soriaga L, Saels V, Herrera-Malaver B, Merlevede A, Roncoroni M, Voordeckers K, Miraglia L, Teiling C, Steffy B, Taylor M, Schwartz A, Richardson T, White C, Baele G, Maere S, Verstrepen KJ. 2016. Domestication and Divergence of *Saccharomyces cerevisiae* Beer Yeasts. Cell 166:1397–1410 e1316.

55. Gibson BR, Storgårds E, Krogerus K, Vidgren V. 2013. Comparative physiology and fermentation performance of Saaz and Frohberg lager yeast strains and the parental species Saccharomyces eubayanus. Yeast 30:255–266.

56. Salazar AN, Gorter de Vries AR, van den Broek M, Brouwers N, de la Torre Cortès P, Kuijpers NGA, Daran J-MG, Abeel T. 2019. Nanopore sequencing and comparative genome analysis confirm lager-brewing yeasts originated from a single hybridization. bioRxiv doi: 10.1101/603480:603480.

57. Lei H, Zhao H, Yu Z, Zhao M. 2012. Effects of wort gravity and nitrogen level on fermentation performance of brewer’s yeast and the formation of flavor volatiles. Appl Biochem Biotechnol 166:1562–1574.

58. Papapetridis I, Verhoeven MD, Wiersma SJ, Goudriaan M, van Maris AJA, Pronk JT. 2018. Laboratory evolution for forced glucose-xylose co-consumption enables identification of mutations that improve mixed-sugar fermentation by xylose-fermenting Saccharomyces cerevisiae. FEMS Yeast Research 18.

59. Wisselink HW, Toirkens MJ, Wu Q, Pronk JT, van Maris AJA. 2009. Novel evolutionary engineering approach for accelerated utilization of glucose, xylose, and arabinose mixtures by engineered *Saccharomyces cerevisiae* strains. Appl Environ Microbiol 75:907–914.

60. Bolat I, Romagnoli G, Zhu F, Pronk JT, Daran JM. 2013. Functional analysis and transcriptional regulation of two orthologs of ARO10, encoding broad-substrate-specificity 2-oxo-acid decarboxylases, in the brewing yeast Saccharomyces pastorianus CBS1483. FEMS Yeast Res 13:505–517.

61. Verduyn C, Postma E, Scheffers WA, Van Dijken JP. 1992. Effect of benzoic acid on metabolic fluxes in yeasts: a continuous-culture study on the regulation of respiration and alcoholic fermentation. Yeast 8:501–517.

62. Pronk JT. 2002. Auxotrophic Yeast Strains in Fundamental and Applied Research. Applied and Environmental Microbiology 68:2095.

63. Solis-Escalante D, Kuijpers NG, Nadine B, Bolat I, Bosman L, Pronk JT, Daran J-MG, Pascale D-L. 2013. amdSYM, a new dominant recyclable marker cassette for *Saccharomyces cerevisiae*. FEMS Yeast Res 13:126–139.

64. Marques WL, Mans R, Marella ER, Cordeiro RL, van den Broek M, Daran J-MG, Pronk JT, Gombert AK, van Maris AJA. 2017. Elimination of sucrose transport and hydrolysis in *Saccharomyces cerevisiae:* a platform strain for engineering sucrose metabolism. FEMS Yeast Res 17:fox006.

65. de Kok S, Yilmaz D, Suir E, Pronk JT, Daran J-MG, van Maris AJA. 2011. Increasing free-energy (ATP) conservation in maltose-grown *Saccharomyces cerevisiae* by expression of a heterologous maltose phosphorylase. Metab Eng 13:518–526.

66. Lee ME, DeLoache WC, Cervantes B, Dueber JE. 2015. A Highly Characterized Yeast Toolkit for Modular, Multipart Assembly. ACS Synth Biol 4:975–986.

67. Gao Y, Zhao Y. 2014. Self-processing of ribozyme-flanked RNAs into guide RNAs in vitro and in vivo for CRISPR-mediated genome editing. Journal of integrative plant biology 56:343–349.

68. Juergens H, Varela JA, Gorter de Vries AR, Perli T, Gast VJ, Gyurchev NY, Rajkumar AS, Mans R, Pronk JT, Morrissey JP. 2018. Genome editing in Kluyveromyces and Ogataea yeasts using a broad-host-range Cas9/gRNA co-expression plasmid. FEMS yeast research 18:foy012.

69. Engler C, Kandzia R, Marillonnet S. 2008. A One Pot, One Step, Precision Cloning Method with High Throughput Capability. PLOS ONE 3:e3647.

70. van Rossum HM, Kozak BU, Niemeijer MS, Duine HJ, Luttik MA, Boer VM, Kötter P, Daran J-MG, van Maris AJA, Pronk JT. 2016. Alternative reactions at the interface of glycolysis and citric acid cycle in *Saccharomyces cerevisiae*. FEMS Yeast Res 16.

71. Gietz RD, Schiestl RH. 2007. High-efficiency yeast transformation using the LiAc/SS carrier DNA/PEG method. Nature protocols 2:31.

72. Pengelly RJ, Wheals AE. 2013. Rapid identification of *Saccharomyces eubayanus* and its hybrids. FEMS Yeast Res 13:156–161.

73. Koren S, Walenz BP, Berlin K, Miller JR, Bergman NH, Phillippy AM. 2017. Canu: scalable and accurate long-read assembly via adaptive k-mer weighting and repeat separation. Genome Res:gr. 215087.215116.

74. Walker BJ, Abeel T, Shea T, Priest M, Abouelliel A, Sakthikumar S, Cuomo CA, Zeng Q, Wortman J, Young SK. 2014. Pilon: an integrated tool for comprehensive microbial variant detection and genome assembly improvement. PloS one 9:e112963.

75. Li H, Durbin R. 2010. Fast and accurate long-read alignment with Burrows–Wheeler transform. Bioinformatics 26:589–595.

76. Holt C, Yandell M. 2011. MAKER2: an annotation pipeline and genome-database management tool for second-generation genome projects. BMC bioinformatics 12:491.

77. Korf I. 2004. Gene finding in novel genomes. BMC bioinformatics 5:59.

78. Stanke M, Keller O, Gunduz I, Hayes A, Waack S, Morgenstern B. 2006. AUGUSTUS: ab initio prediction of alternative transcripts. Nucleic acids research 34:W435–W439.

79. Camacho C, Coulouris G, Avagyan V, Ma N, Papadopoulos J, Bealer K, Madden TL. 2009. BLAST+: architecture and applications. BMC bioinformatics 10:421.

80. Tai SL, Boer VM, Daran-Lapujade P, Walsh MC, de Winde JH, Daran J-M, Pronk JT. 2005. Two-dimensional transcriptome analysis in chemostat cultures combinatorial effects of oxygen availability and macronutrient limitation in Saccharomyces cerevisiae. Journal of Biological Chemistry 280:437–447.

81. Dobin A, Gingeras TR. 2016. Optimizing RNA-Seq mapping with STAR, p 245–262, Data Mining Techniques for the Life Sciences. Springer.

82. Anders S, Pyl PT, Huber W. 2015. HTSeq—a Python framework to work with high-throughput sequencing data. Bioinformatics 31:166–169.

83. Robinson MD, McCarthy DJ, Smyth GK. 2010. edgeR: a Bioconductor package for differential expression analysis of digital gene expression data. Bioinformatics 26:139–140.

84. McCarthy DJ, Chen Y, Smyth GK. 2012. Differential expression analysis of multifactor RNA-Seq experiments with respect to biological variation. Nucleic acids research 40:4288–4297.

85. Anders S, McCarthy DJ, Chen Y, Okoniewski M, Smyth GK, Huber W, Robinson MD. 2013. Count-based differential expression analysis of RNA sequencing data using R and Bioconductor. Nature protocols 8:1765.

86. Entian K-D, Kötter P. 2007. 25 Yeast genetic strain and plasmid collections. Methods in Microbiology 36:629–666.

