## Supplementary material for "Maltotriose consumption by hybrid *Saccharomyces pastorianus* is heterotic and results from regulatory cross-talk between parental sub-genomes": Supplemntary figures with captions

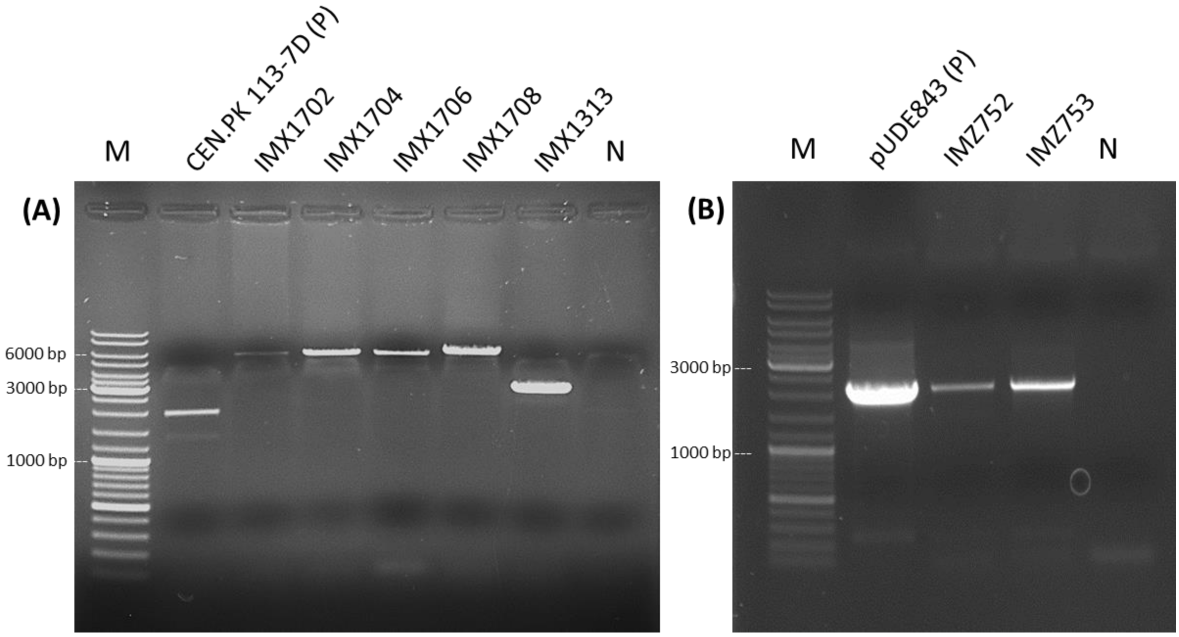


**Supplementary Figure 1 Verification PCR of IMX1702-IMX1708, IMZ752 and IMZ753. (A)** Outside-outside insert PCR amplification of the SGA1 locus or integrated fragments was done using primer pair 4226/4224 . As positive control CEN.PK 113-7D was used. **(B)** Colony PCR to verify the presence of pUDE843 and pUDE844 in IMX1313Δ with primer pair 14454/14455. Gels (TAE 1% agarose) were run at 100V for25 minutes. As marker (M) GeneRuler DNA Ladder Mix (Thermo Scientific) was used.

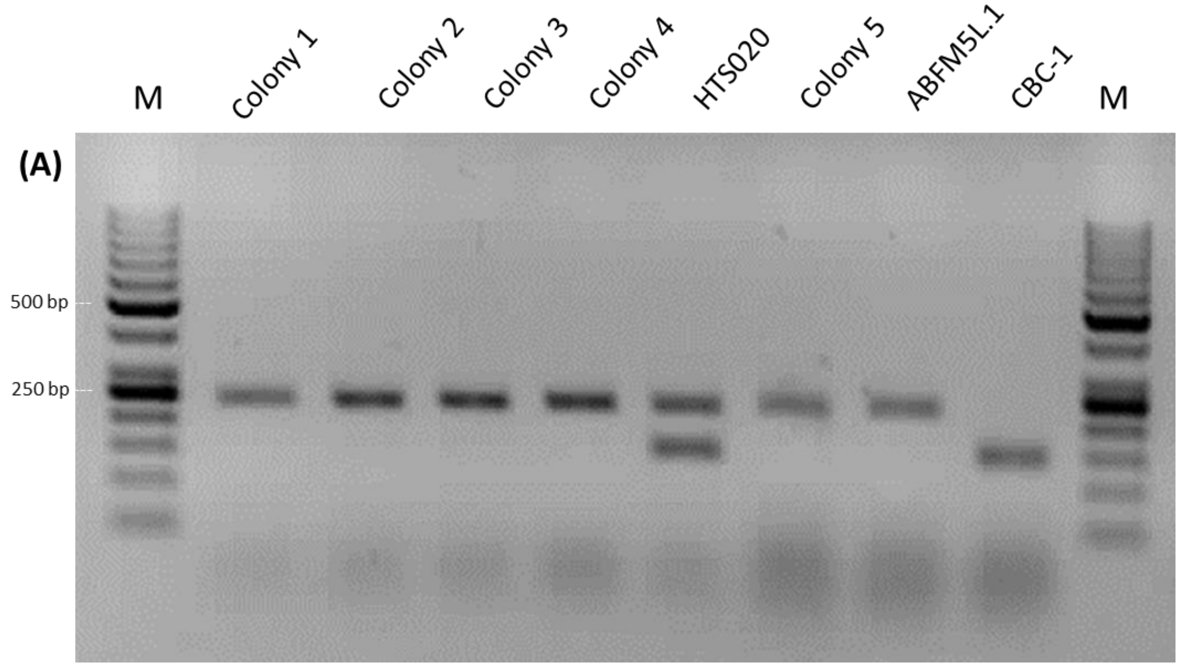


**Supplementary Figure 2 Verification of successful hybridization by multiplex PCR.** Multiplex PCR using primer pairs 8570/8571 (*S. cerevisiae* specific) and 8572/8573 (*S. eubayanus* specific). Lanes contain single colony isolates and ABFM5L.1 and CBC-1 as controls. Gels (TBE 2% agarose) were run at 120V for 40 minutes. As marker (M) GeneRuler 50 bp DNA Ladder (Thermo Scientific) was used.

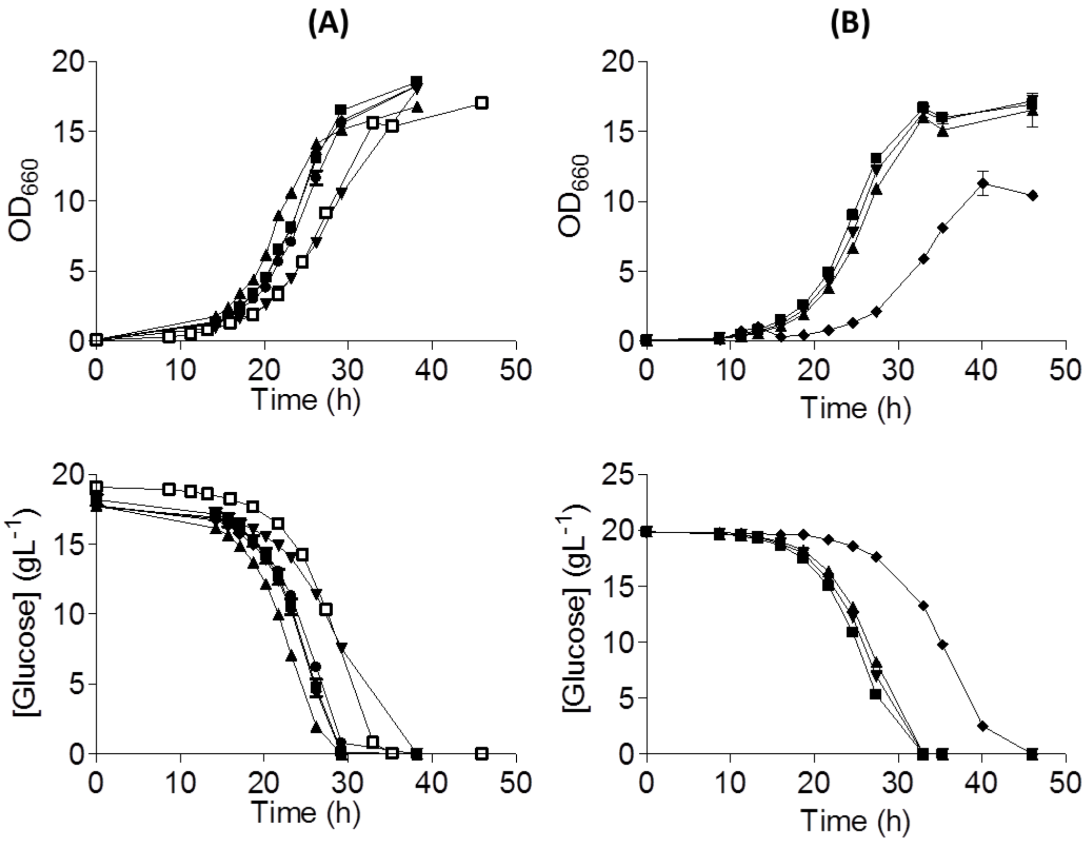


**Supplementary Figure 3 Overexpressing and knockout strains grown on SM glucose. (A)** IMZ616 (■), IMX1365 overexpressing *ScMAL11* (▲), IMX1702 overexpressing *SeMALT1* (▼), IMX1704 overexpressing *SeMALT2* (◆) IMX 1706 overexpressing *SeMALT3* (●) and IMX 1708 overexpressing SeAGT1 (□) were grown on SM 2% glucose at 20 °C. Growth was monitored based on optical density ($OD_{660nm}$) and glucose concentration in culture supernatant was measured by HPLC. Data are presented as average and standard deviation of two biological replicates. **(B)** *S. eubayanus* strains IMK820 (■), IMK823 (▲), IMX1939 (▼) and IMX1940 (◆) were characterized on SM glucose at 20 °C. Growth was monitored based on optical density ($OD_{660nm}$) and glucose concentration in culture supernatant was measured by HPLC. Data are presented as average and standard deviation of two biological replicates.

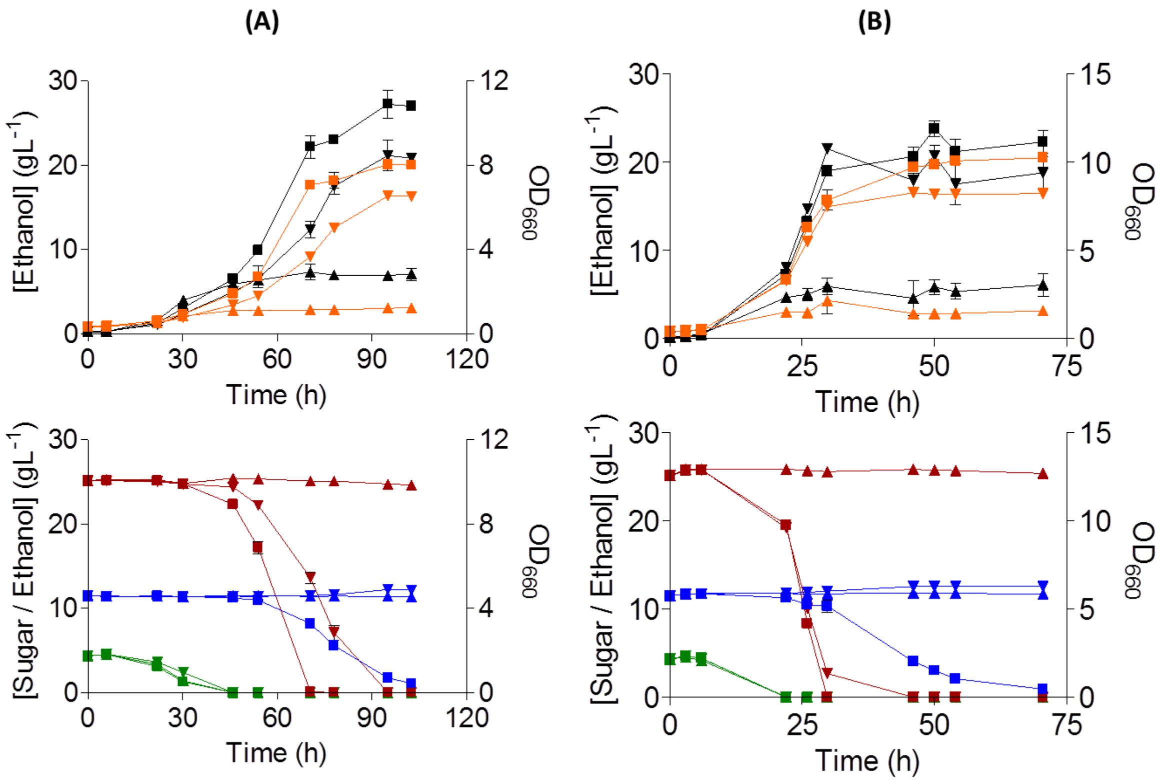


**Supplementary Figure 4 Hybridization of maltotriose deficient *S. cerevisiae* and *S. eubayanus* leading to crosstalk restoring maltotriose utilization, explains *S. pastorianus* phenotype.** Characterization of *S. cerevisiae* CBC-1 (▼), *S.* eubayanus CDFM21L.1 (▲) and hybrid HTS020 (■) on mock wort at 12 °C **(A)** and 20 °C **(B)**. Growth was monitored based on $OD_{660nm}$ (black). Consumption of glucose (green), maltose (red), maltotriose (blue) and production of ethanol (orange) was measured from supernatant by HPLC. Data represents average and standard deviation from biological triplicates.

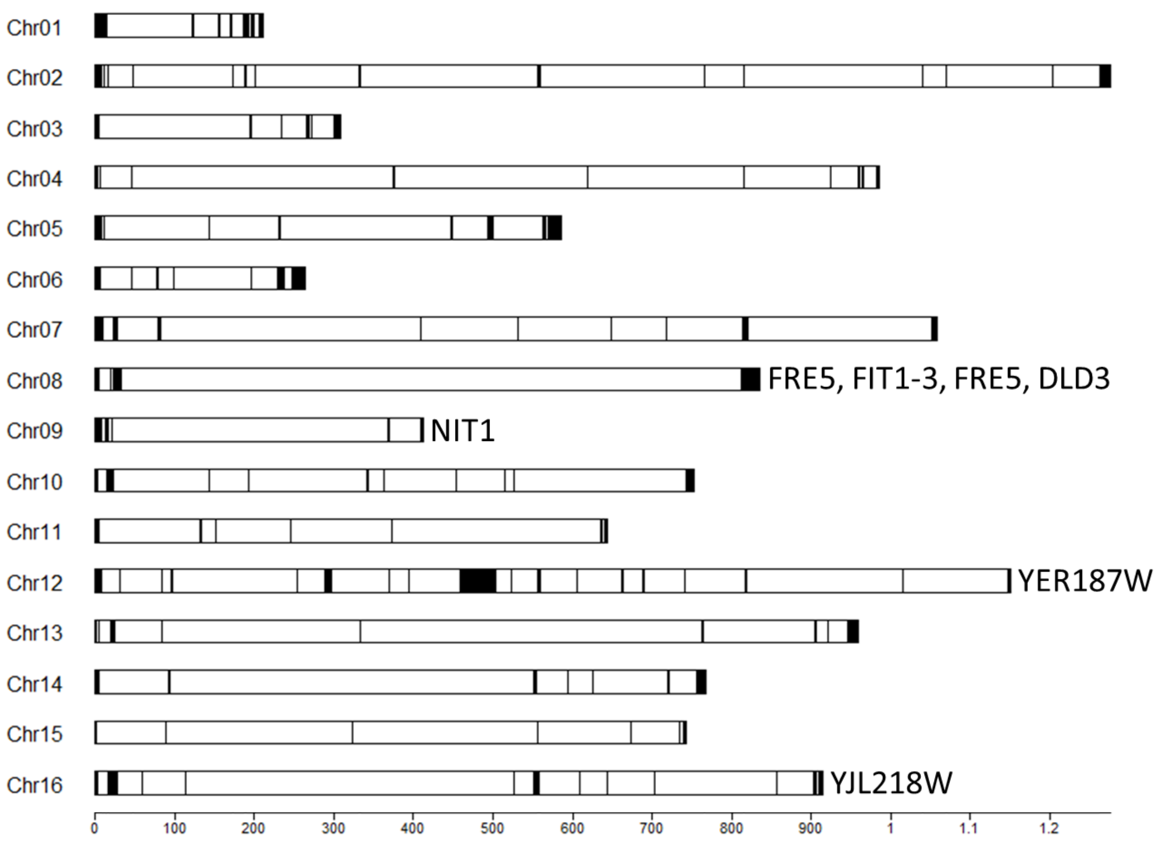


**Supplementary Figure 5 Unmapped and unique regions in CBS 12357$^{T}$**. Per chromosome is indicated which genes are absent in CDFM21L.1.

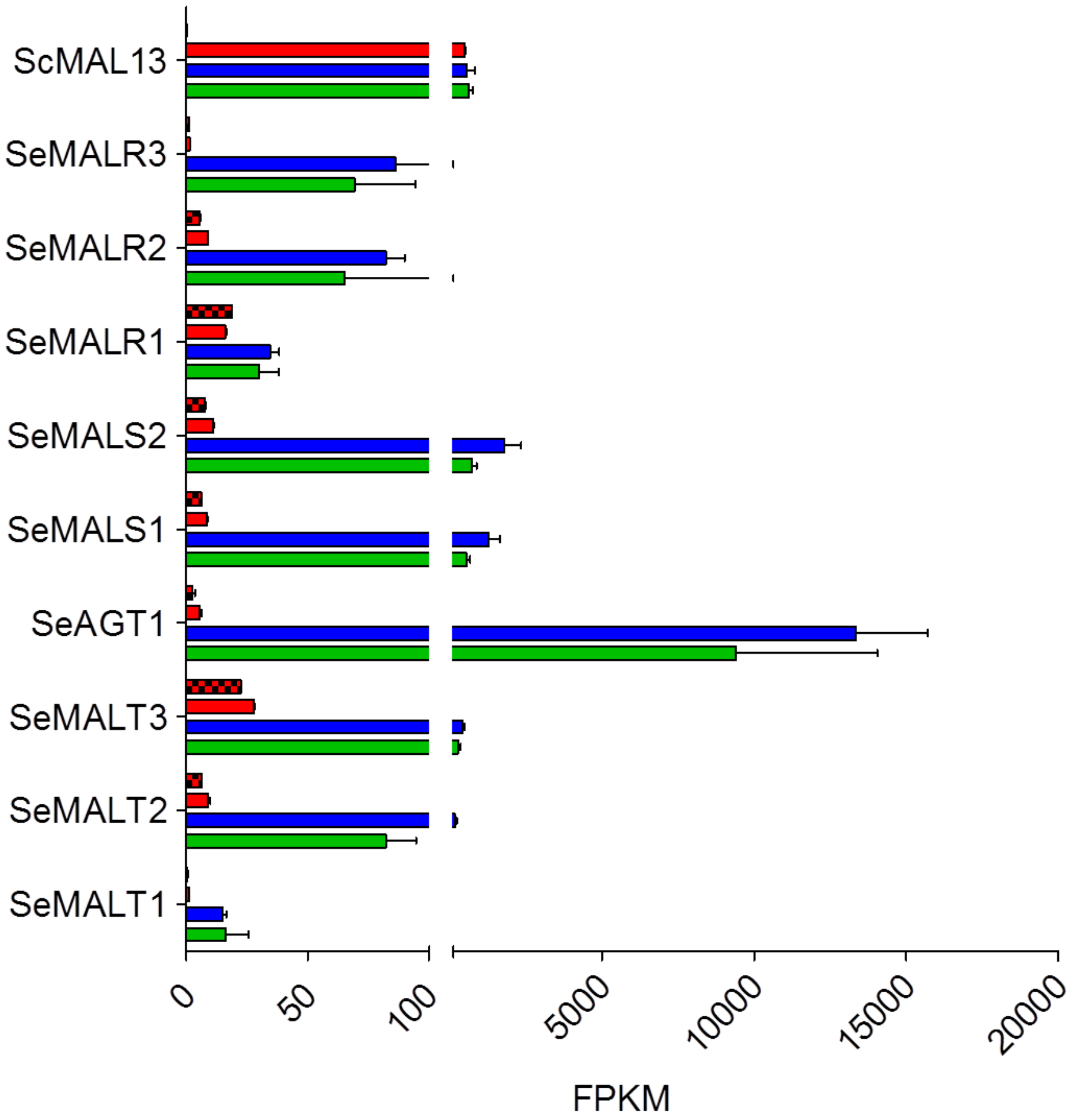


**Supplementary Figure 6 *MAL* gene transcriptomic expression levels.** RNA sequencing data normalized as FPKM of IMX1765 on glucose (red), maltose (blue) and maltotriose (green) and CDFM21L.1 on glucose (red with black squares).
